# Omega-3 Fatty Acid Synergy with Glucocorticoid in Lupus Macrophages: Targeting Pathogenic Pathways to Reduce Steroid Dependence

**DOI:** 10.1101/2025.06.12.658906

**Authors:** Lauren K. Heine, Rance Nault, Jalen Jackson, Ashley N. Anderson, Jack R. Harkema, Andrew J. Olive, James J. Pestka, Olivia F. McDonald

**Affiliations:** Department of Pharmacology and Toxicology, Michigan State University, East Lansing, MI, United States; Institute for Integrative Toxicology, Michigan State University, East Lansing, MI, United States; Department of Microbiology, Genetics, and Immunology, Michigan State University, East Lansing, MI, United States; Department of Pathobiology and Diagnostic Investigation, Michigan State University, East Lansing, MI, United States; Department of Food Science and Human Nutrition, Michigan State University, East Lansing, MI, United States

**Keywords:** fetal liver-derived alveolar-like macrophage (FLAM), glucocorticoid (GC), omega-3 fatty acid, autoimmunity, lupus, interferon (IFN)

## Abstract

**Introduction:** Systemic lupus erythematosus (SLE) is a complex autoimmune disorder characterized by aberrant inflammation, type I IFN-stimulated gene (ISG) expression, and autoantibody production. Glucocorticoids (GCs) like dexamethasone (DEX) are standard long-term SLE treatments but cause significant side effects, highlighting the need for safer steroid-sparing options. Preclinical and clinical studies suggest that dietary supplementation with omega-3 fatty acids (O3FAs), particularly docosahexaenoic acid (DHA), suppress inflammation and autoimmunity associated with SLE disease progression. We explored the steroid-sparing potential of DHA to influence suppressive effects of DEX on pathogenic gene expression.

**Methods:** Macrophages from SLE-prone NZBWF1 mice were first subjected to DHA (5, 10, or 25 µM), DEX (1, 10, 100, or 1000 nM), or DHA+DEX cotreatment. Following pretreatment, cells were exposed to lipopolysaccharide (LPS; 20 ng/mL) to model SLE hyperinflammation. Effects on gene expression were analyzed by qRT-PCR and RNA-seq.

**Results:** qRT-PCR indicated that subinhibitory concentrations of DHA (5-10 µM) potentiated the efficacy of low-dose DEX (1-100 nM) in suppressing LPS-induced ISG expression (e.g., *Irf7*, *Oasl1*, *Rsad2*), amplifying the effects of DEX monotherapy by 10- to 100-fold. SynergyFinder analysis confirmed that DHA and DEX interacted synergistically in suppressing ISG expression, with significant inhibition observed at concentrations as low as 1 nM DEX and 5 µM DHA. RNA-seq revealed that combining suboptimal DHA (10 μM) and DEX (100 nM) induced 247 differentially expressed genes (DEGs) at 4 hr and 347 DEGs at 8 hr post-LPS, dramatically surpassing the effects of each treatment alone. Functional enrichment analysis indicated that DHA+DEX cotreatment robustly suppressed immune and inflammatory pathways while promoting proliferative and metabolic processes, reflecting a shift from inflammatory (M1) to pro-resolving (M2) macrophage phenotypes. DHA and DEX countered LPS effects by i) downregulating common transcription factors (TFs) canonically associated with inflammation (e.g., NF-κB, AP-1, STATs, and IRF1), ii) upregulating shared regulatory factors involved in inflammation resolution (e.g., YBX1, EGR1, and BCL6), and iii) selectively influencing other regulatory factors.

**Discussion:** Altogether, DHA and DEX synergistically suppress inflammation by targeting common and unique molecular pathways in SLE macrophages, favoring the pro-resolving M2 phenotype. O3FA-GC cotreatment might facilitate reducing requisite steroid dosages for SLE management.

## 1 Introduction

Systemic lupus erythematosus (SLE, lupus) is a debilitating autoimmune disease characterized by chronic inflammation, aberrant type I IFN-stimulated gene (ISG) responses, loss of self-tolerance, and multi-organ damage driven by genetic susceptibility and environmental triggers (1, 2). Genome-wide association studies in patients with SLE have identified more than 150 risk loci that converge on pathways regulating IFN signaling and immune cell activation (1, 3). Environmental exposures, such as airborne pollutants, infections, and ultraviolet light, amplify genetic predispositions to SLE by triggering oxidative stress and aberrant nucleic acid sensing (4, 5).

Macrophage hyperactivity plays a central role in SLE pathogenesis (6, 7) by perpetuating tissue injury through dysregulated cytokine production, phagocytic dysfunction, and sustained type I IFN secretion—a hallmark of SLE observed in 60-80% of patients (8, 9). Macrophage hyperactivity is mediated by the activation of pattern-recognition receptors such as toll-like receptors (TLRs), which detect both pathogen-associated molecular patterns (PAMPs) and damage-associated molecular patterns (DAMPs) (6, 7), or by activation of cytokine receptors specific for type I IFN, TNFα, IL-1, or IL-6 (8, 10, 11). This is exemplified preclinically in SLE-prone NZBWF1 mouse alveolar macrophages, which exhibit heightened pathogenic gene expression following activation of TLR4 by PAMPs like lipopolysaccharide (LPS) or DAMPs unleashed by toxic silica particles (5, 12, 13). TLR4 activation triggers MAPK/NF-κκB signaling and IFN regulatory factors (IRFs), resulting in the increased expression of proinflammatory and type I IFN-stimulated genes (ISGs) (14, 15). This cascade creates a feed-forward loop that enhances antigen presentation, promotes autoantibody production by plasma cells, and activates autoreactive T cells (16, 17). Accordingly, airborne environmental triggers, such as LPS and silica, accelerate the onset and progression of SLE in NZBWF1 mice, highlighting the crucial role of AM hyperactivity in lupus pathogenesis.

Glucocorticoids (GCs; steroids) remain the cornerstone of lupus treatment, as they suppress key inflammatory pathways, including NF-κB and MAPK signaling (18, 19). However, chronic GC use at moderate-to-high doses is associated with severe adverse side effects, including osteoporosis, hyperglycemia, muscle atrophy, cardiovascular complications, and increased infection risk (20–22). Consistent with this notion, we found that while moderate-to-high dose GC treatment of silica-exposed NZBWF1 mice at translationally relevant dosages inhibits proinflammatory and autoimmune gene expression, it also elicits significant muscle wasting and hyperglycemia without improving survival outcomes (23). These findings highlight the critical need for steroid-sparing adjunctive therapies that effectively control inflammation while minimizing patient health risks.

Emerging evidence positions marine oil-derived omega-3 fatty acids (O3FAs), such as docosahexaenoic acid (DHA), as promising anti-inflammatory agents for adjunctive treatment in systemic lupus erythematosus (SLE) and other autoimmune diseases (24–26). Mechanistically, DHA exerts its effects through multiple pathways: i) altered receptor function due to lipid raft composition (27) and size (28), ii) activating anti-inflammatory TFs such as PPARγ (29), iii) inhibiting NF-κB (30), iv) disrupting cholesterol synthesis, v) upregulating NFE2L2 (NRF2)-associated genes (31), and vi) producing specialized pro-resolving mediators like resolvins and maresins (32). We recently reported that, in a cohort of 418 participants with SLE, higher serum levels of O3FAs, particularly DHA, were associated with favorable outcomes, including reduced SLE scores, less pain, and improved sleep quality (33). In preclinical studies of silica-triggered SLE in NZBWF1 mice, we demonstrated that dietary DHA supplementation suppresses IFN-stimulated and proinflammatory gene expression and consequent pulmonary inflammation and lupus nephritis (5, 34–36).

Given their potential for complementary anti-inflammatory mechanisms, we posit that O3FAs could be used as adjuncts to reduce GC dosages needed to suppress SLE progression. To test this hypothesis, we preclinically modeled SLE hyperinflammation using LPS activation of novel self-renewing fetal liver-derived alveolar-like macrophages (FLAMs) derived from NZBWF1 mice (37). The results showed that combining subinhibitory concentrations of DHA with low-dose dexamethasone (DEX) creates a potent synergy that robustly suppresses IFN-stimulated and proinflammatory gene expression induced by LPS in the SLE macrophages. Cotreatment outperformed individual treatments by targeting key pathways and transcription factors (TFs) involved in inflammation and resolution. These findings support the idea that O3FA-GC cotreatment may be a feasible steroid-sparing strategy for SLE management.

## 2 Methods

### 2.1 Self-renewing fetal liver-derived SLE macrophages

The Institutional Animal Care and Use Committee (IACUC) at Michigan State University (MSU; AUF# 201800113) approved all animal experimental protocols for this study. SLE-prone NZBWF1 mice (Jackson Laboratories, Bar Harbor, ME) were housed at MSU’s animal facility, which was maintained at a constant temperature (21-24°C), humidity (40-55%), and 12-hr light/dark cycle. After the mice were bred, the dams were euthanized between gestational days 14 and 18. Mice were euthanized via CO_2_ inhalation for 10 minutes to ensure death to neonates, and cervical dislocation was used as a secondary form of euthanasia for the dam. Fetuses were promptly removed from the dam, and the loss of access to the maternal blood supply served as the secondary form of euthanasia for the neonates. Fetal livers were excised from neonates and further processed to generate fetal liver-derived alveolar-like macrophage (FLAM) cell cultures as previously described (31, 37). Briefly, fetal livers were dissociated in sterile phosphate-buffered saline (PBS) to create a single-cell suspension. Suspensions were filtered through a 70-µm filter and centrifuged at 220 *x g* for 5 minutes. Two wash steps were performed using sterile PBS before resuspending cells in modified RPMI media (mRPMI, Thermo Fisher), which contained 10% fetal bovine serum (FBS, Thermo Fisher), 1% penicillin-streptomycin (P/S, Thermo Fisher), 30 ng/mL murine granulocyte-monocyte colony-stimulating factor (GM-CSF, PeproTech), and 20 ng/mL recombinant human TGF-β1 (PeproTech). Cells were plated in 10 cm culture plates (one liver per plate). mRPMI media was replaced every ∼2 days until cells created an adherent monolayer and exhibited a round AM-like morphology (∼1 wk). Cells were then frozen in liquid nitrogen until needed for this study. FLAMs were thawed and cultured in mRPMI media for this experiment, and cells between passages 10-11 were used for the following studies.

### 2.2 Study 1. Quantitative RT-PCR analysis of DHA and DEX cotreatment effects on LPS-induced ISG expression

#### 2.2.1 Experimental design

FLAMs were seeded in 12-well plates at -48 hr in mRPMI media. At -24 hr, cells were gently washed with PBS, and media was replaced with mRPMI containing 0.25% FBS and 0, 5, 10, or 25 µM DHA (NuChek Prep, Elysian, MN). At -1 hr, FLAMs were treated with mRPMI media containing 0.25% FBS and 0, 1, 10, 100, or 1000 nM DEX (Sigma-Aldrich). At 0 hr, FLAMs were treated with mRPMI medium containing vehicle (VEH/CON) or containing 20 ng/mL LPS (LPS/VEH; *Salmonella enterica* serotype Typhimurium containing <1% impurities, Millipore Sigma). Cells were collected at 4 and 8 hr for qRT-PCR.

#### 2.2.2 qRT-PCR

Total cellular RNA was extracted using RNeasy Mini Kits (Qiagen) according to the manufacturer’s instructions. RNA was eluted using RNase-free water provided by the RNeasy kit and quantified using the Nanodrop ND-1000 spectrophotometer (Thermo Fisher Scientific, Waltham, MA). cDNA was prepared from RNA using a High-Capacity cDNA Reverse Transcriptase Kit (Thermo Fisher Scientific). Taqman assays were performed in technical triplicate using the Takara Bio Smart Chip real-time PCR system, with assistance from the MSU Genomics Core, to assess gene expression. Expression of ISGs (*Mx1, Irf7, Ifit1, Isg15, Oasl1, and Rsad2*) and housekeeping genes (*Actb and Hprt*) was assessed. ΔCt values were calculated by subtracting the average raw Ct value of both housekeeping genes from the raw Ct value for each gene of interest. ΔΔCt values were calculated by subtracting the average ΔCt value of the respective VEH/CON group from the average ΔCt value of the LPS/VEH treatment. ΔΔCt values are shown in units of fold increase relative to LPS/VEH for each gene of interest. Similarly, to assess DHA/DEX treatment on LPS-stimulated FLAMs, ΔΔCt values were calculated by subtracting the average ΔCt value of the respective LPS/VEH group from the average ΔCt value of the corresponding DHA/DEX treatment. ΔΔCt values are shown in units of fold increase relative to LPS/VEH for each gene of interest.

### 2.3 Study 2. RNA-seq and functional analysis of DHA and DEX cotreatment effects on immune pathways

#### 2.3.1 Experimental Design

FLAMs were seeded in 6-well plates at -48 hr in mRPMI media. At -24 hr, cells were gently washed with PBS, and media was replaced with mRPMI media containing 0.25% FBS with or without 10 µM DHA. At -1 hr, FLAMs were treated with mRPMI media containing 0 or 100 nM DEX. At 0 hr, FLAMs were treated with mRPMI medium (VEH/CON) or media containing 20 ng/mL LPS. Culture cohorts were collected for fatty acid analysis (4 hr), gene expression by RNA-seq (4 and 8 hr), and cytokine secretion by ELISA (24 hr). Treatment groups were as follows: (i) VEH/CON, (ii) LPS/VEH, (iii) DHA/LPS, (iv) DEX/LPS, and (v) DHA+DEX/LPS.

#### 2.3.2 Fatty acid analysis

Cell pellets were preserved in 100% methanol at -80°C before fatty acid composition analysis using gas chromatography with flame ionization detection at OmegaQuant (Sioux Falls, SD). The procedure involved transferring pellets to screw-cap glass vials and adding an internal standard, di-C23:0 PL. A modified Folch extraction was performed, followed by thin-layer chromatography (TLC) separation using a solvent mixture of hexane, ethyl ether, and acetic acid (8:2:0.15). The phospholipid band from the TLC plate was collected and treated with methanol containing 14% boron trifluoride. HPLC-grade water and hexane were added after heating at 100°C for 10 minutes. The mixture was vortexed and centrifuged for phase separation. The hexane layer underwent gas chromatography analysis using a GC2010 Gas Chromatograph with a specific capillary column. Fatty acids were quantified by comparison with a standard mixture and an internal standard. Di-C23:0 PL was used to calculate recovery efficiency. The analysis identified 24 different saturated fatty acids (SFAs), monounsaturated fatty acids (MUFAs), omega-6 fatty acids (O6FAs), and omega-3 fatty acids (O3FAs). Results were expressed as a percentage of total identified fatty acids.

#### 2.3.3 RNA-seq

Cells were lysed using RLT lysis buffer (Qiagen), and RNA was isolated from cells using RNeasy Mini Kits (Qiagen). RNA was quantified with Qubit (Thermo Fisher Scientific), and integrity was verified with TapeStation (Agilent Technologies). Samples (RNA integrity numbers >8) were library-prepped at the MSU Genomics Core using the Illumina Stranded mRNA Library Preparation, Ligation Kit with IDT for Illumina Unique Dual Index adapters following the manufacturer’s recommendations, except that half-volume reactions were used. Libraries were pooled in equimolar proportions and quantified using the Invitrogen Collibri Quantification qPCR kit. Samples were sequenced on the NovaSeq 6000 S4 flow cell in a 2x150bp paired-end format using a NovaSeq v1.5, 300-cycle reagent kit. Base calling was performed using Illumina Real-Time Analysis (RTA) v3.4.4, and the RTA output was demultiplexed and converted to FastQ format with Illumina Bcl2fastq v2.20.0. Following quality control using FastQC, reads were aligned to the mouse reference genome (GRCm39, release 107) using STAR (version 2.3.7a) (38). Normalization and differential expression analysis were performed using DESeq2 (39) in R (version 4.1.2). Genes were considered differentially expressed when | fold change | ≥ 1.5 and the adjusted p-value ≤ 0.05.

#### 2.3.4 Functional analysis

Gene set enrichment analysis was performed using the fgsea package in R on gene expression datasets ranked by fold-change and gene sets from the Gene Set Knowledgebase (GSKB) (40) filtered only to include Gene Ontology (GO) and KEGG gene sets (41). The pathway-level information extractor (PLIER) tool (42) was used to summarize gene expression signatures for all treatment conditions, except for unstimulated controls, using the same gene sets as prior knowledge. Differences in latent variable (LV) estimates between conditions were determined by three-way ANOVA with DHA treatment, DEX treatment, and time as factors.

#### 2.3.5 Transcription factor (TF) analysis

TF analysis was performed using the decoupleR package (43). DESeq2 differential expression analyses sorted by fold-change were used as input with the DoRothEA collection of TF-gene interactions with a level of confidence “A – highest confidence” (44).

#### 2.3.6 Enzyme-linked immunosorbent assay

Representative ISG-related proteins (i.e., IFN-β, CCL2, CXCL10) that were identified with qRT-PCR and RNA-seq were measured in supernatants of treated FLAMs by enzyme-linked immunosorbent assay (ELISA). Specifically, IFN-β was measured using a LumiKine™ Xpress mIFN-β 2.0 kit (InvivoGen, San Diego, CA), and other proteins were measured using corresponding DuoSet ELISA kits (R&D Systems, Minneapolis, MN) according to the manufacturer’s instructions.

### 2.4 Data visualization and statistics

GraphPad Prism Version 10 (GraphPad Software, La Jolla, California, USA, www.graphpad.com) was used to visualize fatty acid, quantitative RT-PCR, and cytokine data. These data were subjected to the ROUT outlier test (Q = 1%) and then the Shapiro-Wilk test (p < 0.01) to identify outliers and assess normality, respectively. Data failing to meet the assumption for normality were analyzed using the non-parametric Kruskal-Wallis test, followed by Dunn’s *post-hoc* test. Data that met assumptions for normality and equal variances were analyzed by parametric one-way analysis of variance (ANOVA) followed by Tukey’s *post-hoc* test. Data are shown as mean ± standard error of the mean, with p < 0.05 considered statistically significant. The SynergyFinder 3.14.0 R package (45) was used to evaluate inhibitory interactions between DHA and DEX on ISG expression. Synergy scores were calculated using the zero-interaction potency (ZIP) model (46). Visualizations of RNA-seq differential expression and functional enrichment analyses were generated using GraphPad Prism and R.

## 3 Results

### 3.1 Study 1: DHA and DEX interact synergistically to inhibit ISG expression in SLE FLAMs

LPS stimulation in NZBWF1 FLAMs induced robust upregulation of representative ISGs, including *Irf7, Mx1, Ifit1*, *Isg15*, *Oasl1*, and *Rsad2*. DHA monotherapy concentration-dependently suppressed this IFN response. Significant ISG suppression was observed at 25 µM (p < 0.05 vs. LPS control) **(Figure 1A)**, while lower concentrations (5 to 10 µM) showed a significant decrease (**Figure 1B, C**). DEX monotherapy at 1 µM effectively suppressed ISG expression, whereas lower doses (≤100 nM) demonstrated incomplete and variable transcriptional inhibition **(Figure 1A-C).** Combining subinhibitory DHA (5 to 10 µM) with low-dose DEX (1 to 100 nM) enhanced the suppression of LPS-driven ISGs (p < 0.05 vs. individual treatments), exceeding additive effects **(Figure 1B, C).** This combination amplified DEX’s potency by 10- to 100-fold, achieving near-complete transcriptional silencing. This potentiation was observed across multiple ISGs, indicating broad modulation of type I IFN-regulated genes and signaling pathways.

**Figure 1.**
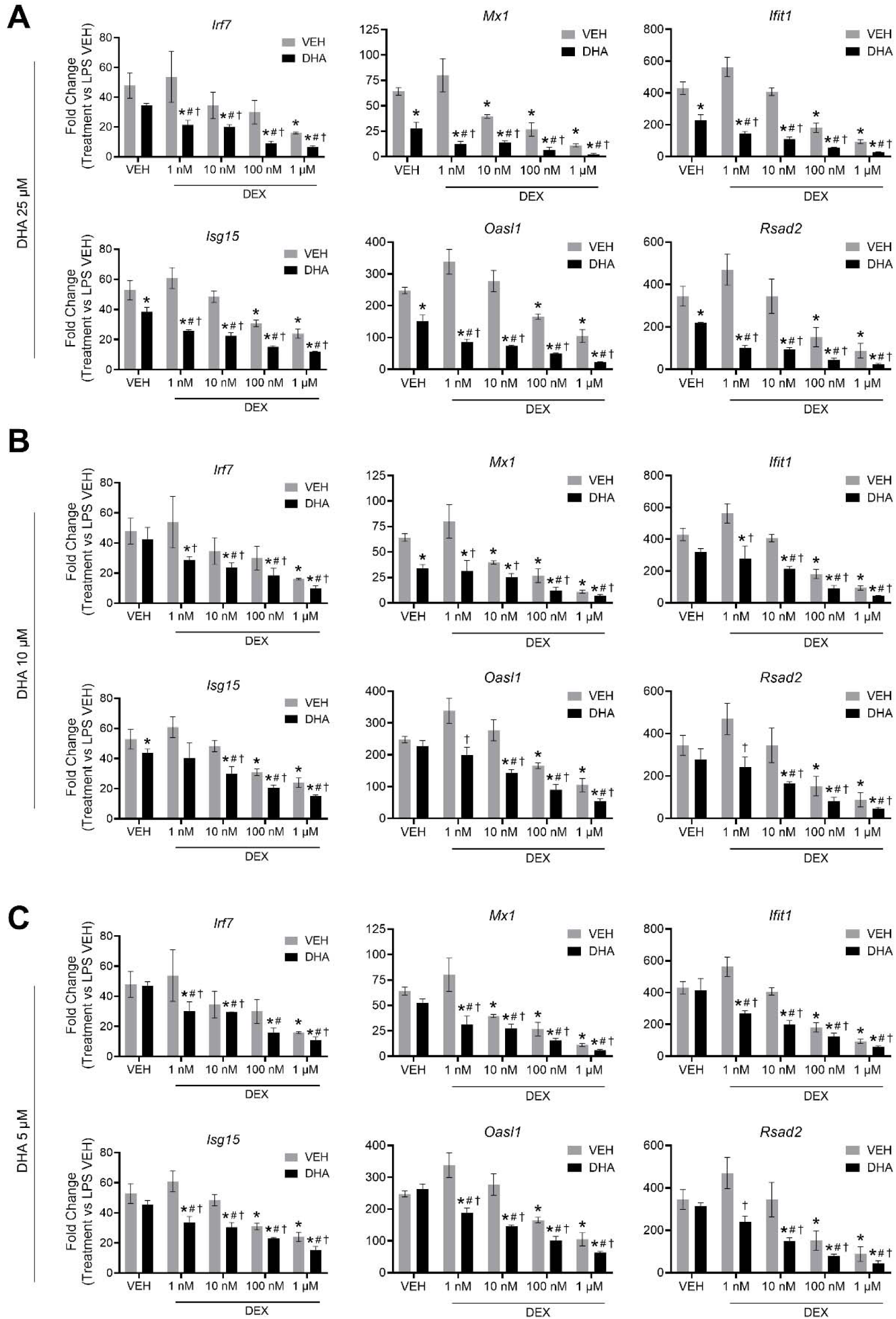
DHA and DEX monotherapy or cotreatment suppress IFN-stimulated gene expression. Cells were pretreated with either VEH containing no DHA or RPMI media containing 25 µM **(A)**, 10 µM **(B)**, or 5 µM **(C)** DHA at -24 hr. Cells were then treated with VEH containing no DEX or varying concentrations of DEX (1 nM to 1 µM), -1 hr prior to LPS treatment. qRT-PCR was performed on FLAMs stimulated with LPS (20 ng/mL) for 4 hr. Fold change is shown as DHA and/or DEX treatment relative to LPS VEH ± SEM. n=3 biological replicates. p<0.05; *Significant compared to LPS VEH; #Significant compared to DHA alone; †Significant compared to DEX alone.

Using SynergyFinder 3.14.0 and the ZIP synergy model, we quantified the synergy between DHA and DEX related to the inhibition of ISG expression. At all concentrations, DHA and DEX demonstrated significant synergistic interactions in inhibiting *Irf7* **(Figure 2A),** *Oasl1* **(Figure 2B)**, *Rsad2* **(Figure 2C)**, *Ifit1* **(Supplementary** Figure 1A), *Isg15* **(Supplementary** Figure 1B), and *Mx1* **(Supplementary** Figure 1C). Synergy was most robust when FLAMs were pretreated with 1 nM or 10 nM DEX in conjunction with 5 µM DHA **(Figure 2** and **Supplemental Figure 1).** At higher concentrations of DHA (i.e., 10 µM and 25 µM) and DEX (i.e., 100 nM and 1000 nM), ZIP synergy scores were still greater than 0, indicating a smaller magnitude of synergy, as the pathways were more robustly inhibited by monotherapies at these higher concentrations.

**Figure 2.**
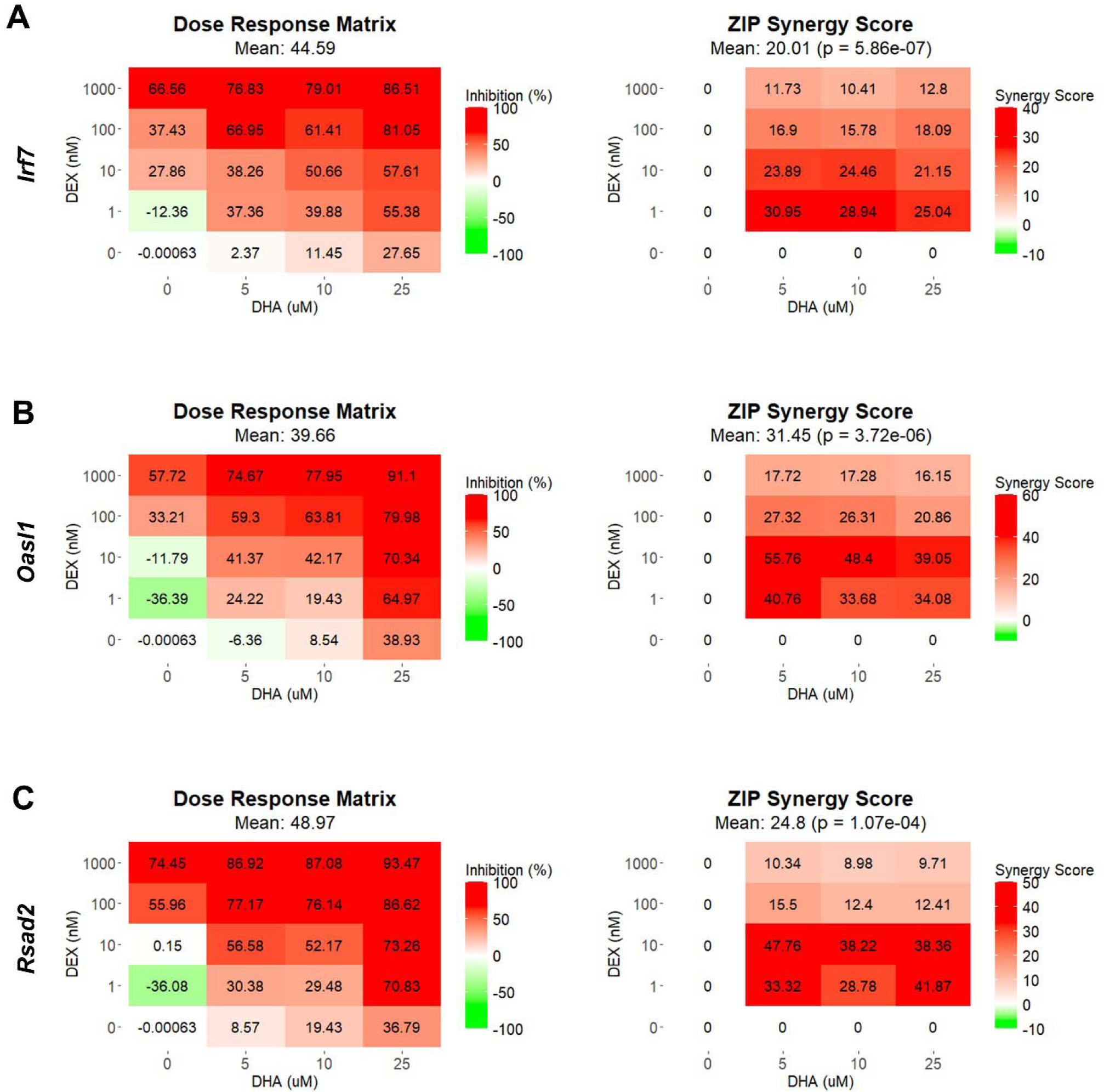
DHA and DEX synergistically inhibit the expression of IFN-stimulated genes. *Irf7* **(A),** *Oasl1* **(B),** and *Rsad2* **(C)** were measured by qRT-PCR in FLAMs stimulated with LPS (20 ng/mL) for 4 hr. Cells were pretreated with either VEH containing no DHA or RPMI media containing 25 µM, 10 µM, or 5 µM DHA at -24 hr. Cells were then treated with VEH containing no DEX or varying concentrations of DEX (1 nM-1 µM) -1 hr prior to LPS treatment. SynergyFinder version 3.14.0 generated inhibition matrices and ZIP synergy matrices for each gene. Inhibition matrices show the average of 3 experimental replicates. Individual and mean ZIP synergy scores were calculated using an average of 3 experimental replicates. Synergy score > 0, synergistic interaction; synergy score = 0, additive effect; synergy score < 0, antagonistic interaction.

Consistent with gene expression, LPS exposure stimulated secretion of IFN-β and selected ISG products (CCL2 and CXCL10) after 24 hr compared to VEH-treated control cells **(Supplemental Figure 2**). These responses were significantly attenuated by DHA and DEX monotherapies. When DHA and DEX were administered in combination, the secretion of IFN-β, CCL2, and CXCL10 was further reduced. Accordingly, combining DHA and DEX enhanced suppression of ISG protein expression, further underscoring their potential as a combined therapeutic strategy for modulating TLR4-driven pathogenic gene responses.

### 3.2 Study 2. RNA-seq and functional analysis of DHA and DEX cotreatment effects on immune pathways in SLE FLAMs

#### 3.2.1 Treatment with DHA but not DEX skews cellular phospholipid profiles

Fatty acid profiles of phospholipids were profoundly altered by DHA treatment (**Figure 3A, B**). DHA, the primary O3FA, rose from 4.9% in the VEH/CON group to 12.0% and 11.9% in the LPS/DHA and LPS/DHA+DEX groups, respectively, with total O3FA levels reaching 18.0% and 17.8%, compared to 9.5% in the VEH/CON group. In tandem with these observations, O6FAs decreased in DHA-treated groups. Linoleic acid (C18:2ω6) fell from 1.1% in the VEH/CON group to 0.8% in the LPS/DHA+DEX group, while arachidonic acid (C20:4ω6) decreased from 8.3% to 6.6% and 6.3% in the LPS/DHA and LPS/DHA+DEX groups, respectively. Likewise, MUFAs declined significantly with DHA treatment. Oleic acid (C18:1ω9), the dominant MUFA, dropped from 36.1% in the VEH/CON group to 22.2% and 21.4% in the LPS/DHA and LPS/DHA+DEX groups, respectively. SFAs increased notably in the LPS/DHA and LPS/DHA+DEX groups compared to the VEH/CON group with palmitic acid (C16:0), the main SFA, rising from 18.9% in the VEH/CON group to 28.5% and 25.0% in the LPS/DHA and LPS/DHA+DEX groups, respectively. Similarly, stearic acid (C18:0) increased to 14.0% in both groups, compared to the VEH/CON group. Overall, DHA supplementation remodeled fatty acid profiles in the phospholipid fraction by increasing the levels of O3FA and SFA while reducing those of MUFA and O6FA. Adding DEX slightly enhanced some trends but did not significantly alter the effects of DHA.

**Figure 3.**
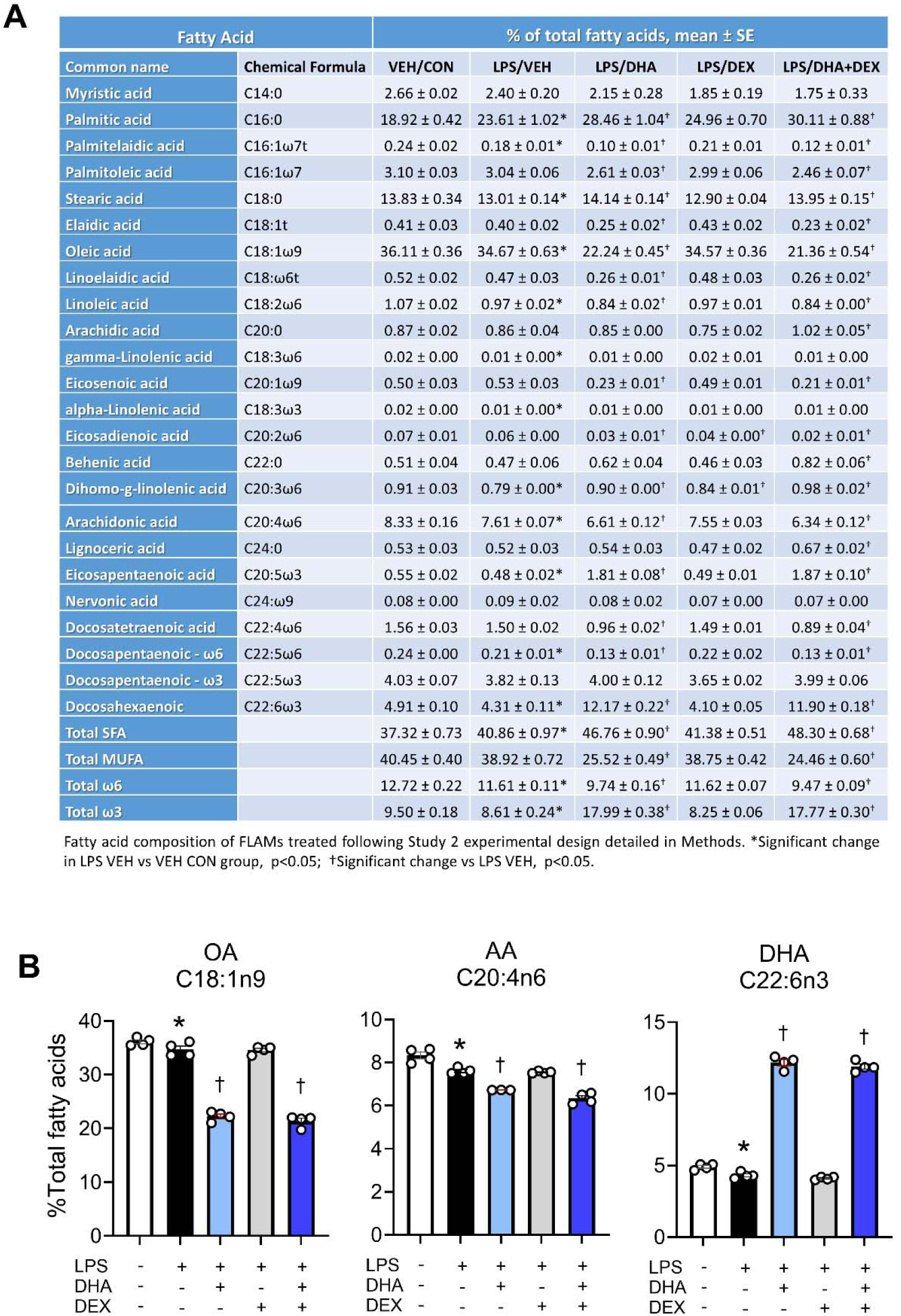
Treatment with DHA but not DEX skews cellular phospholipid profiles. **(A)** FLAMs were treated with DEX, DHA, and/or LPS as described and analyzed for major fatty acids as described in the Methods section. **(B)** DHA supplementation resulted in increased phospholipid DHA with accompanied decreases in arachidonic acid (AA) and oleic acid OA. *Significant differences (p<0.05) between VEH/CON and LPS/VEH were determined using Student’s t-test; †Significant differences (p<0.05) between LPS/VEH and LPS/DHA, LPS/DEX, or LPS/DHA+DEX treatments were determined using one-way ANOVA.

#### 3.2.2 LPS treatment significantly alters the transcriptome, enriching pathways related to inflammation and the immune response

RNA-seq analysis revealed that LPS treatment resulted in 3,632 significantly differentially expressed genes (DEGs) at 4 hr and 3,571 DEGs at 8 hr compared to VEH/CON-treated FLAMs **(Figure 4A)**. Of these DEGs, 1860 were upregulated at 4 hr and 1998 at 8 hr. Conversely, 1772 DEGs were downregulated at 4 hr and 1573 at 8 hr. These responses highlight the dynamic nature of gene regulation during LPS-induced inflammation.

**Figure 4.**
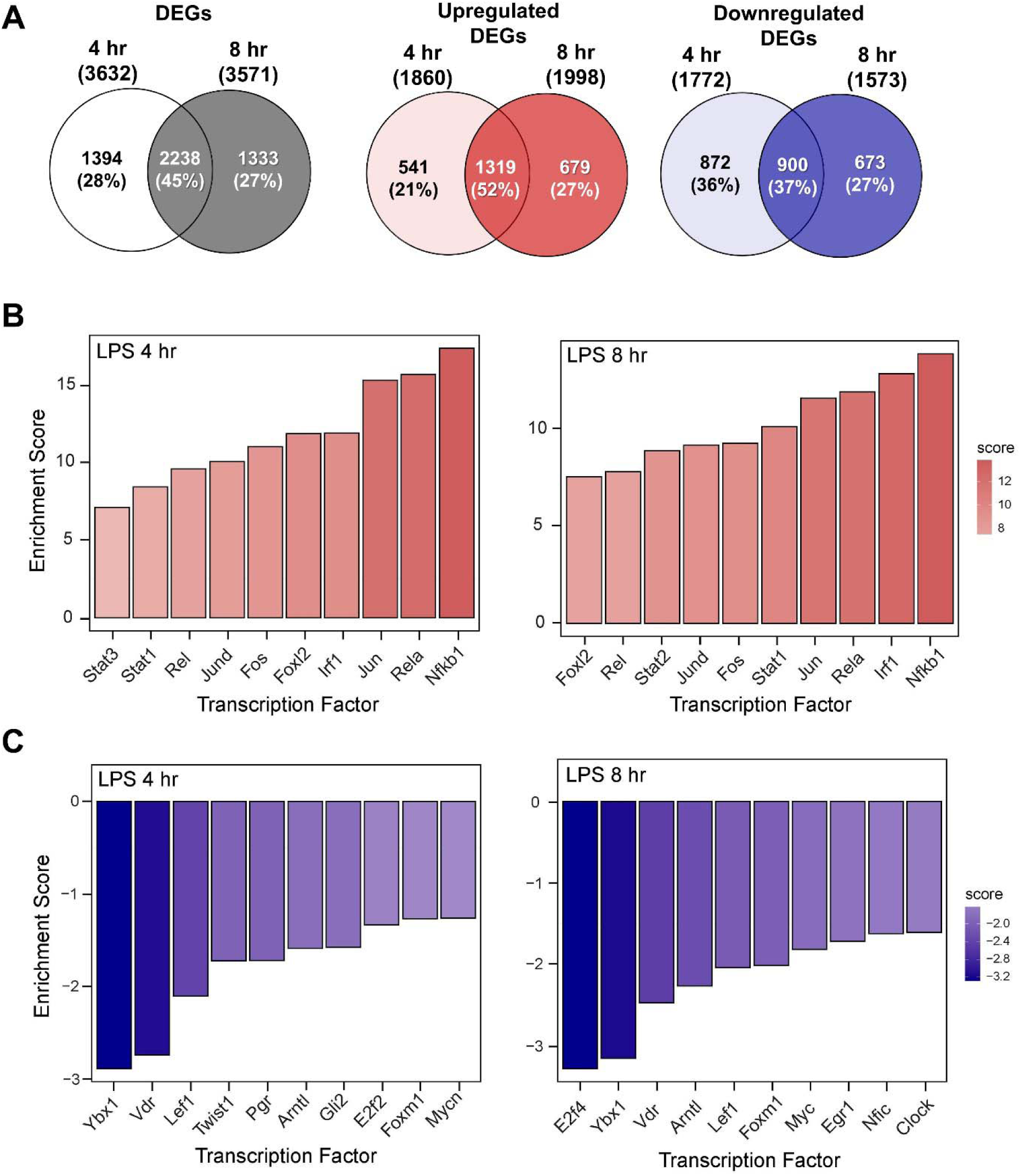
LPS induces proinflammatory transcriptional responses in FLAMs. **(A)** Differentially expressed genes (DEGs) were determined using DESeq2 (39) filtering for genes exhibiting a |log2 fold change| >= 2 and adjusted p-value <= 0.05 between the LPS/VEH group and VEH/CON group. Venn diagrams for total DEGs, upregulated DEGs, and downregulated DEGs are shown for the 4-hr time point (left circle), 8-hr time point (right circle), and both time points (intersection). **(B, C)** The top 10 inferred **(B)** active and **(C)** inactive TFs following LPS treatment were identified for both time points using decoupleR (43).

The top 10 inferred active and inactive transcription factors (TFs) responding to LPS treatment were identified using decoupleR for both 4- and 8-hr time points. Enrichment scores were elevated for canonical proinflammatory TFs, including NF-κB (Rel, Nfkb1), AP-1 (Jun, JunD, Fos), STATs (Stat1,2,3), and Irf1 **(Figure 4B)**. Notably, nine out of ten TFs remained among the most positively enriched regulators across both time points, indicating a coordinated regulation of inflammatory responses through immediate-early transcriptional programs. In addition, anti-inflammatory and regulatory transcription factors (TFs), such as Ybx1, Lef1, E2f2/4, Vdr, and Mycn/Myc, were negatively enriched at both time points **(Figure 4C)**. These coordinindicatedTF signatures highlight their potential vital role in modulating LPS-induced immune and inflammatory signaling in FLAMs.

Functional enrichment analysis using GSEA revealed that LPS treatment at both time points induces distinct transcriptional responses, as shown by normalized enrichment scores (NES) across biological processes **(Figure 5)**. Positive NES were evident for immune/inflammatory responses, chemokine/cytokine activity, responses to viruses, LPS, IFN, and double-stranded DNA (dsDNA). Elevated NES values were equivalent or higher at 4 hr than 8 hr, illustrating temporal dynamics of LPS-induced immune activation. Conversely, negative NES were associated with cell division, DNA replication, lipid metabolism, spindle, and microtubule motor activity, consistent with suppression of proliferative and metabolic processes following LPS stimulation.

**Figure 5.**
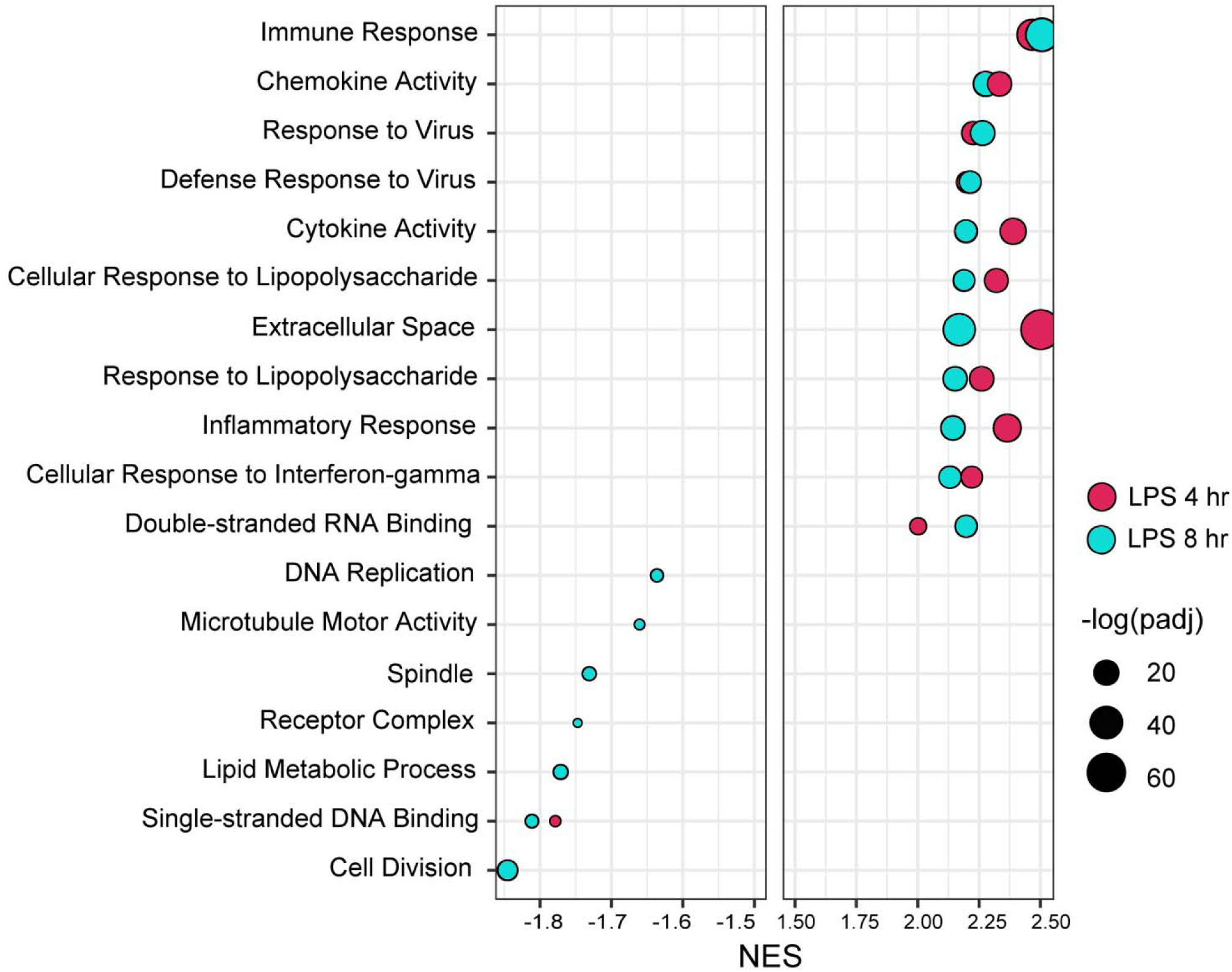
LPS induces inflammation and innate immunity and suppresses proliferative and metabolic biological processes. Gene set enrichment analysis was performed using the fgsea package in R on gene expression datasets ranked by fold-change and gene sets from the Gene Set Knowledgebase (GSKB) (40) filtered only to include Gene Ontology (GO) and KEGG gene sets (41). At both 4-hr and 8-hr time points, LPS induced immune and inflammatory pathways, including cytokine/chemokine signaling and responses to viruses, LPS, IFN, and dsDNA. These immune-related processes showed equal or greater activation at 4 hr compared to 8 hr, highlighting the rapid and dynamic nature of the inflammatory response. Simultaneously, pathways related to cell proliferation and metabolism (blue bars) exhibited significant negative enrichment, including cell division, DNA replication, lipid metabolism, and microtubule-associated processes, indicating coordinated suppression of growth and metabolic functions during LPS-induced inflammation.

#### 3.2.3 DHA+DEX cotreatment suppresses LPS proinflammatory responses in FLAMs

Sub-inhibitory DHA monotherapy (10 μM) in LPS-primed FLAMs resulted in 16 DEGs at 4 hr and 83 DEGs at 8 hr (**Figure 6A**). DEX monotherapy (100 μM) resulted in 48 DEGs at 4 hr and 160 DEGs at 8 hr. There were 0 and 10 common DEGs at 4 and 8 hr shared between DHA and DEX, respectively. Consistent with synergy observed in Study 1, DHA+DEX cotreatment significantly increased DEGs being expressed at 4 hr (247) and 8 hr (347). DHA+DEX cotreatment in LPS-primed FLAMs resulted in 247 DEGs at 4 hr and 347 DEGs at 8 hr. There were 15 and 40 percent overlaps of DEGs between cotreatment and monotherapies at 4 and 8 hr, respectively. Functional enrichment analysis using GSEA indicated strong negative enrichment for many LPS-upregulated gene pathways (**Figure 6B**).

**Figure 6.**
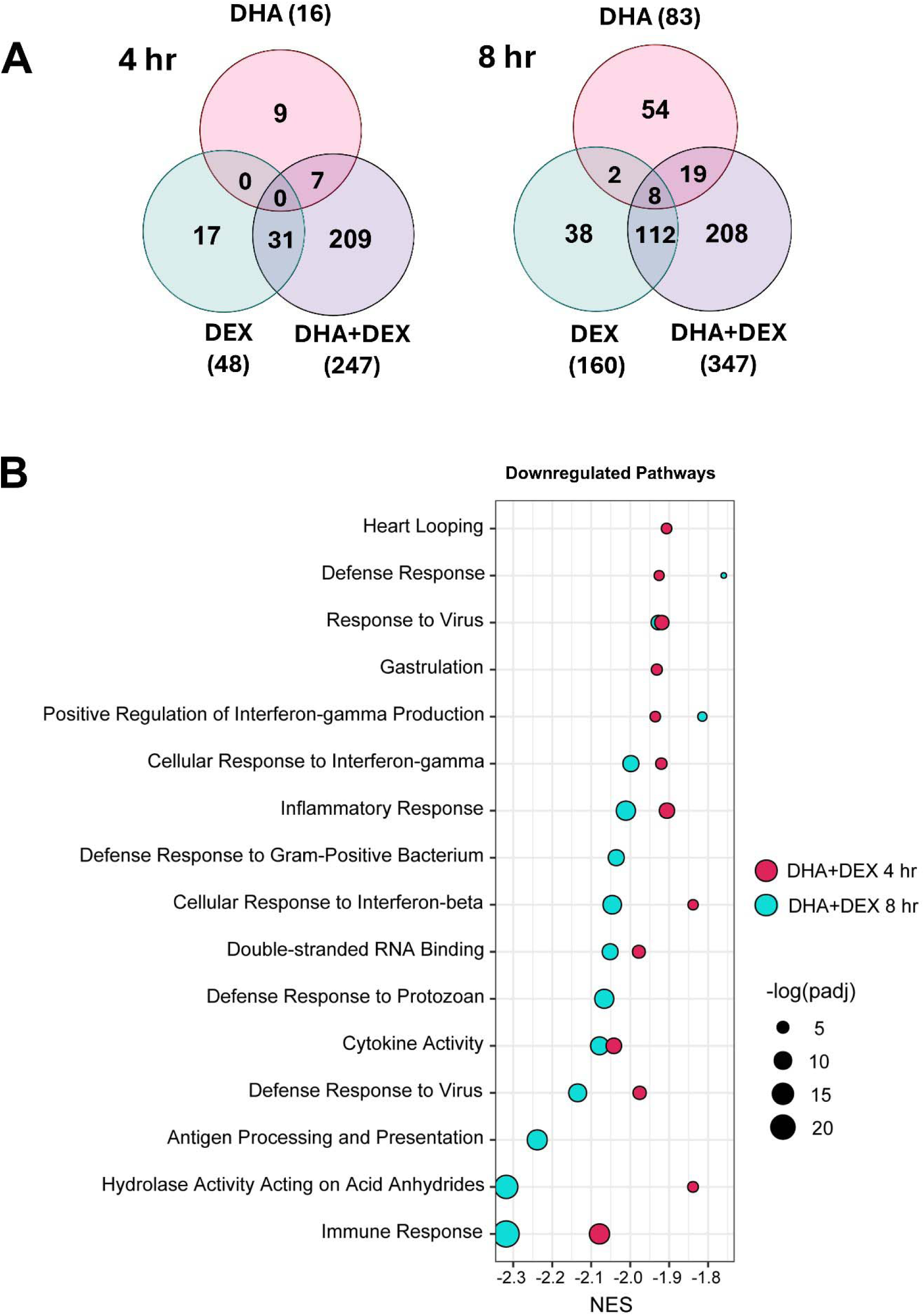
DHA and DEX monotherapies and DHA+DEX cotreatment suppress LPS-stimulated transcriptional responses in FLAMs. (A) Differentially expressed genes (DEGs) were determined using (39) filtering for genes exhibiting a |log2 fold change| >= 2 and adjusted p-value <= 0.05 between DHA, DEX, or DHA+DEX treatment relative to LPS/VEH. Venn diagrams for the number of treatment-dependent unique gene symbols for 4- and 8-hr time points. (B) Functional enrichment analysis using the GSEA method, as described in Figure 5 legend, indicates negative enrichment for LPS-upregulated gene pathways.

Functional enrichment analysis using PLIER was further used to identify biological processes associated with DHA+DEX treatment, revealing that cotreatment significantly altered pathways related to the cellular response to IFN, antigen processing/presentation, and NAD ADP-ribosyl transferase (**Figure 7A, B**). The top 40 genes most significantly altered by DHA+DEX treatment were extracted from the identified pathways and depicted in a heatmap (**Figure 7C**). Genes altered with DEX+DHA treatment were associated with IFN and antiviral response (e.g., *Mx1, Mx2, Oasl1, Ifit1, Sp140, Gm5431*), MHC antigen processing and presentation (e.g., *H2-DMa, H2-Ab1, H2-Aa, H2-Eb1*), cytokine receptor signaling (e.g., *CD74, Ccr5, Ccl2*), and apoptosis and proliferation (e.g., *Mxd1, Daxx, Zeb1*).

**Figure 7.**
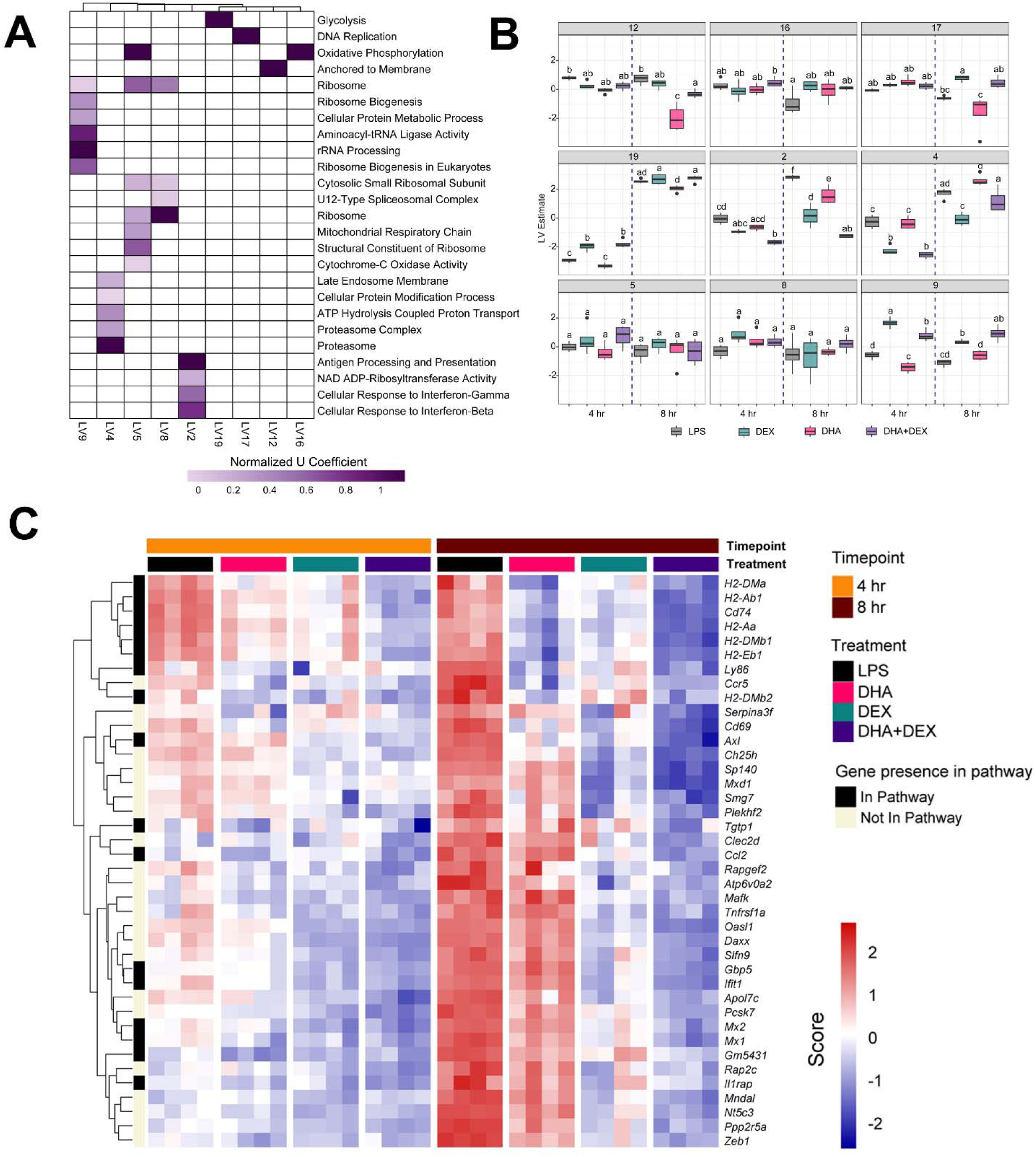
Identification of biological processes enriched by DHA+DEX cotreatment using PLIER analysis. (A) Functional enrichment analysis using the pathway-level information extractor (PLIER) method was used to identify significantly enriched biological processes associated with DHA+DEX treatment. PLIER was used to identify high-confidence latent variables (LVs; AUC >= 0.7 and FDR <= 0.05) mapped to Gene Ontology (GO) and KEGG gene sets. **(B)** LV estimates for high-confidence LVs are shown for each treatment group. Treatment groups were assessed by 3-way ANOVA, and different letters indicate significant differences (p <= 0.05). **(C)** Heat maps of the top 40 genes enriched by cotreatment were determined using PLIER. The color scale corresponds to the scaled expression value, with red being highly expressed genes and blue corresponding to downregulated genes.

Log2-fold changes in representative gene responses resulting from treatment are depicted in **Figure 8**. Individual DEGs pertaining to type I/II IFNs (**Figure 8A**), cytokine signaling (**Figure 8B**), and antigen processing and presentation (**Figure 8C**) were downregulated by DHA and/or DEX compared to LPS treatment at 8 hr. Downregulation of each gene was potentiated with cotreatment, and DHA+DEX treated cells were significantly different compared to cells treated with DHA and DEX individually. Combinatorial effects were observed for some, but not all, genes at the 4-hr time point.

**Figure 8.**
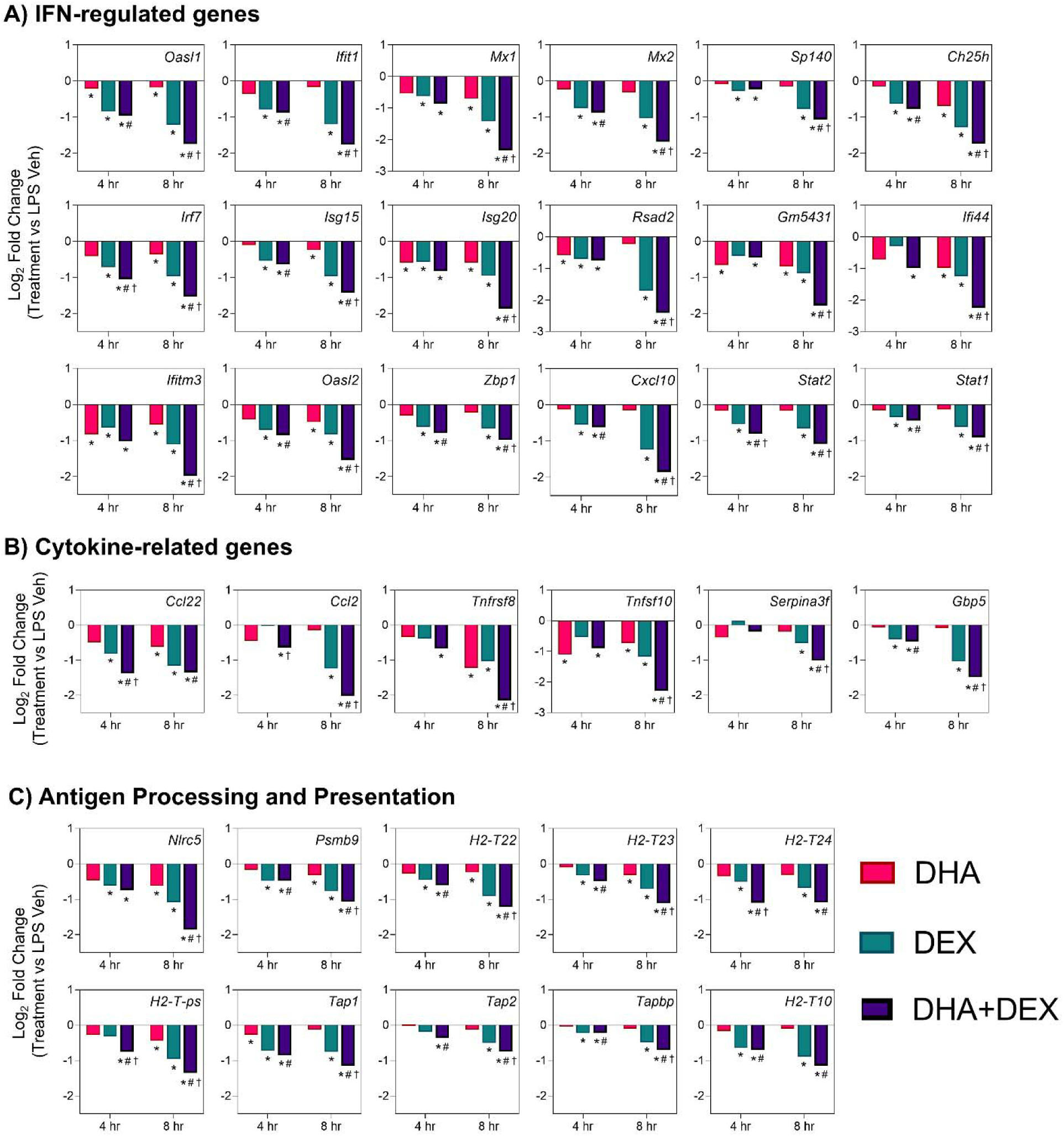
DHA+DEX combinatorial effects on suppression of innate immune response genes. Representative individual differentially expressed genes related to **(A)** type I IFNs, **(B)** cytokine signaling and **(C)** antigen processing and presentation that exhibited combinatorial effects compared to individual DHA or DEX treatment were extracted from the RNA-seq dataset. Log2 fold change was determined relative to LPS/VEH. p<0.05; *Significant compared to LPS/VEH; #Significant compared to DHA monotherapy within respective time point; †Significant compared to DEX monotherapy within respective time point.

#### 3.2.4 DHA, DEX, and DHA+DEX treatments impact transcriptional responses in LPS-primed FLAMs

The effects of DHA, DEX, and DHA+DEX treatment on LPS-induced DEGs and predicted TF activity regulation in LPS-primed cells are depicted in **Figures 9, 10,** and **11**. DHA treatment alone led to the upregulation of 5 DEGs at 4 hr and 13 DEGs at 8 hr with no overlapping DEGs, and the downregulation of 11 DEGs at 4 hr and 100 DEGs at 8 hr with 2 overlapping DEGs (**Figure 9A**). Treatment with DEX alone contributed to the upregulation of 21 DEGs at 4 hr and 41 DEGs at 8 hr, with 3 overlapping DEGs, and the downregulation of 18 DEGs at 4 hr and 110 DEGs at 8 hr, with 6 overlapping DEGs (**Figure 10A**). DHA+DEX combination treatment resulted in the upregulation of 112 DEGs at 4 hr and 48 DEGs at 8 hr, with 8 overlapping DEGs, and the downregulation of 75 DEGs at 4 hr and 239 DEGs at 8 hr, with 52 overlapping DEGs (**Figure 11A**).

**Figure 9.**
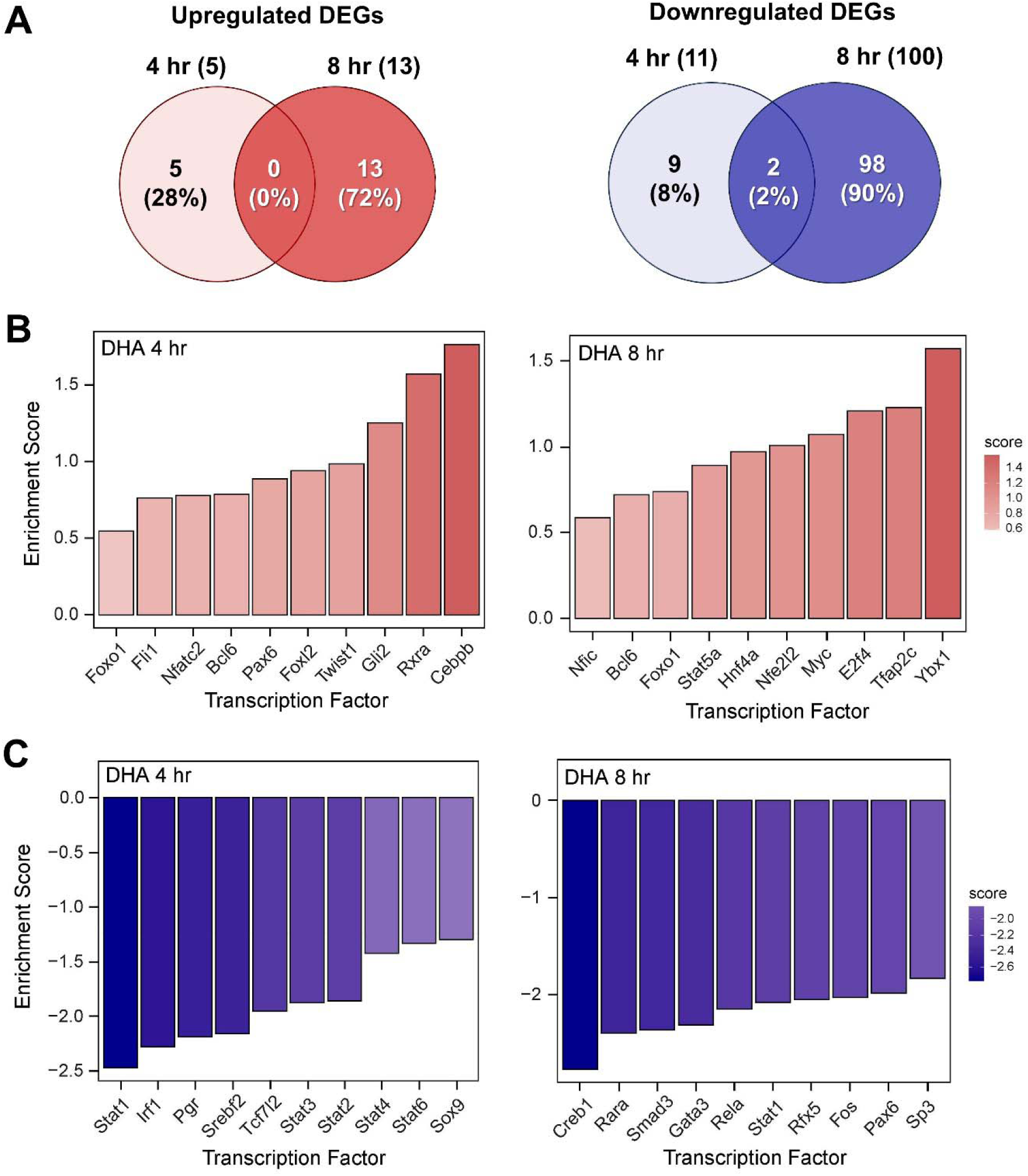
DHA monotherapy influences LPS-induced DEGs and transcription factor regulation. **(A)** Downregulated and upregulated differentially expressed genes (DEGs) were determined using DESeq2 (39) filtering for genes exhibiting a |log2 fold change| >= 2 and adjusted p-value <= 0.05 between the LPS/VEH group and VEH/CON group. Venn diagrams for downregulated DEGs are shown for the 4 hr time point (left circle), 8 hr time point (right circle), and both time points (intersection). The top 10 inferred (B) active and (C) inactive TFs following DHA treatment were identified at 4 hr and 8 hr using decoupleR (43).

**Figure 10.**
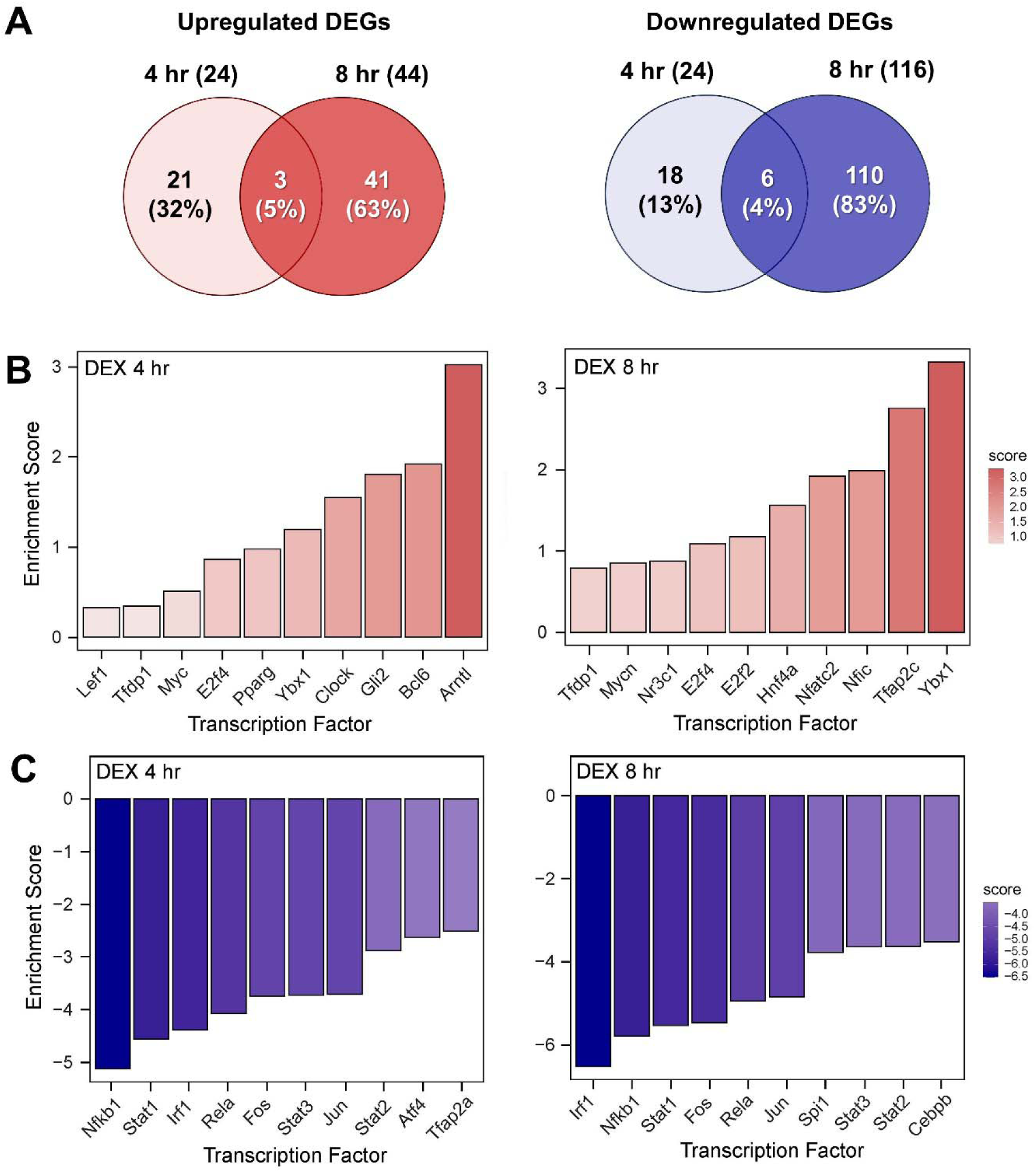
DEX monotherapy influences LPS-induced DEGs and transcription factor regulation. **(A)** Downregulated and upregulated differentially expressed genes (DEGs) were determined using DESeq2 (39) filtering for genes exhibiting a |log2 fold change| >= 2 and adjusted p-value <= 0.05 between the LPS/VEH group and VEH/CON group. Venn diagrams for downregulated DEGs are shown for the 4 hr time point (left circle), 8 hr time point (right circle), and both time points (intersection). The top 10 inferred (B) active and (C) inactive TFs following DHA treatment were identified at 4 hr and 8 hr using decoupleR (43).

**Figure 11.**
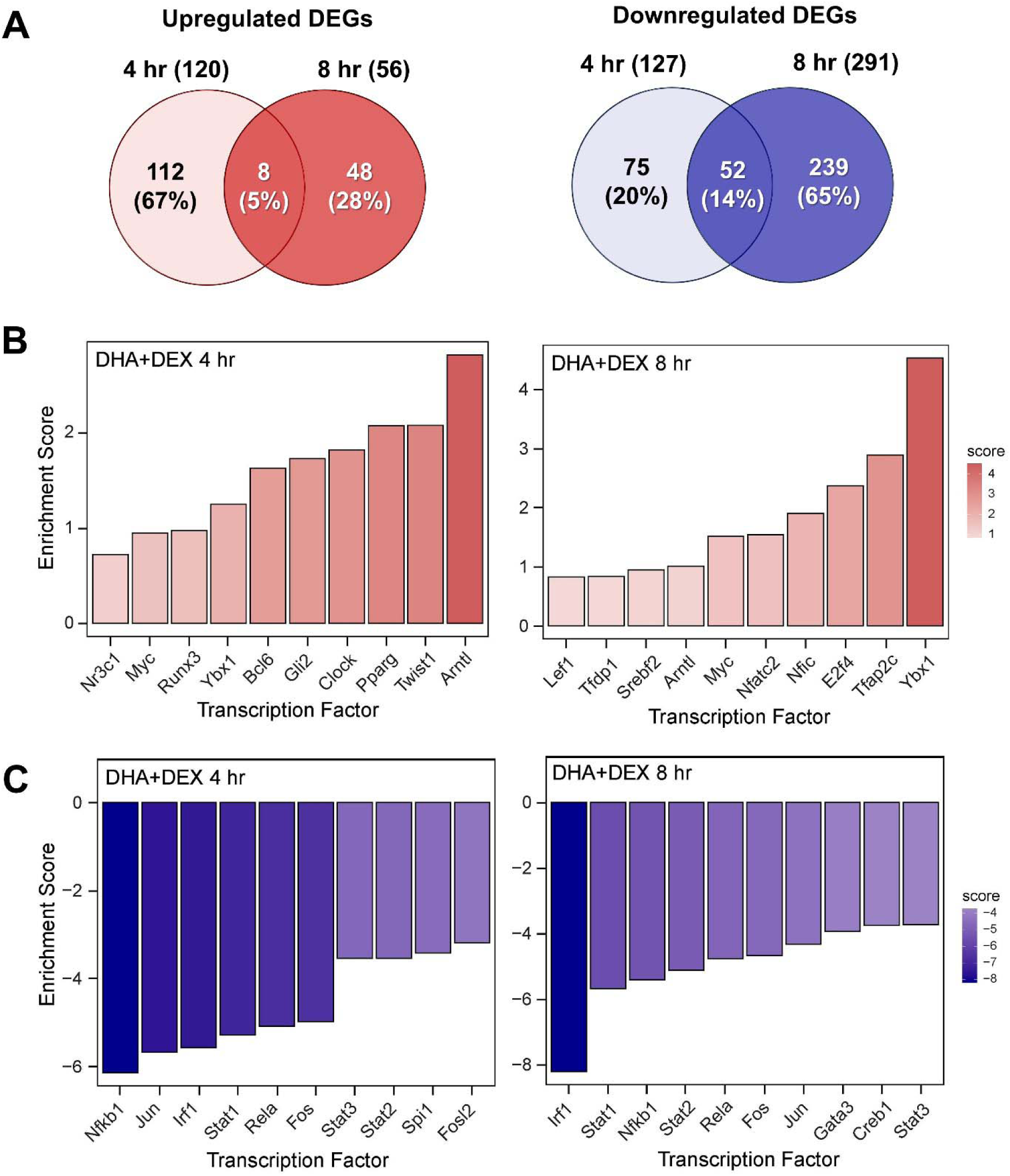
DHA+DEX cotreatment robustly modulates LPS-induced DEGs and transcription factor regulation. **(A)** Downregulated and upregulated differentially expressed genes (DEGs) were determined using DESeq2 (39) filtering for genes exhibiting a |log2 fold change| >= 2 and adjusted p-value <= 0.05 between the LPS/VEH group and VEH/CON group. Venn diagrams for downregulated DEGs are shown for the 4 hr time point (left circle), 8 hr time point (right circle), and both time points (intersection). The top 10 inferred **(B**) active and **(C)** inactive TFs following DHA+DEX cotreatment were identified at 4 hr and 8 hr using decoupleR (43).

DHA **(Figure 9B)**, DEX **(Figure 10B)**, and DHA+DEX **(Figure 11B)** treatments all resulted in positive enrichment of TFs that are involved with reducing inflammation (e.g., Ybx1) (47, 48), proliferation (e.g., Bcl6, E2f4)(49–52), differentiation and development (e.g., Myc, Gli2, Nfic) (53–57), and metabolism (e.g., Arntl) (58, 59). DEX and DHA+DEX treatment led to positive enrichment scores for Pparg, Nr3c1, and Clock, which are involved with fatty acid metabolism, GC signaling, and circadian rhythm regulation, respectively (21, 60, 61). DHA and DHA+DEX contributed to enrichment for Twist1, which is involved with reducing inflammation (62), while DHA alone selectively enriched for anti-inflammatory factors Cebpb, Rxra, FlII, Nfe2l2, Stat5a, and Foxo1 (63–70)(**Figure 9B**).

DHA (**Figure 9C**), DEX (**Figure 10C**), and DHA+DEX (**Figure 11C**) treatment all led to negative enrichment of TFs that regulate proinflammatory cytokines (e.g., Rel, Fos) and ISGs (e.g., Irf1, Stat1, Stat2, Stat3). DHA specifically contributed to the negative enrichment of factors that were not affected by DEX, including Pgr, Srebf2, Tcf7l2, Sox9, Creb1, Rarα, Smad3, Gata3, Rfx5, Pax6, and Sp3 (**Figure 9C**). DEX and DHA+DEX treatment resulted in significant negative enrichment of Nfkb1 and Jun, which are components of the NF-κB and AP-1 heterodimers, respectively (**Figures 10C** and **11C**).

## 4 Discussion

### 4.1 Synopsis

Macrophages orchestrate immune responses through gene expression finely tuned by a complex interplay of TFs to balance inflammation, antimicrobial defense, and resolution. Dysregulation of these pathways can contribute to SLE onset and drive pathogenesis, underscoring the importance of macrophages as therapeutic targets in SLE management. While cellular and molecular mechanisms of GC and O3FA treatments have been individually studied for their anti-inflammatory effects on macrophages (13, 31, 71–73), the combined impact of these treatments on transcriptional networks remains unexplored. We addressed this question by determining how cotreatment with DEX and DHA modulates critical regulatory hubs and the transcriptional landscape in LPS-stimulated NZBWF1 FLAMs. Our findings support the conclusion that DHA+DEX cotreatment synergistically reverses LPS-induced changes in transcription and regulatory factor activities, markedly attenuating expression of IFN-stimulated and proinflammatory genes that contribute to SLE pathogenesis. This synergy reveals novel crosstalk mechanisms between O3FAs and GCs, highlighting the potential therapeutic value of combining lipidomic and pharmacological approaches to combat SLE-associated inflammation.

### 4.2 DHA and DEX suppress LPS activation of canonical proinflammatory regulators

As shown in **Figure 12**, our deconvolution findings closely align with established TLR4 signaling pathways where LPS activates both MyD88-dependent and TRIF-dependent pathways to drive M1 macrophage polarization (74). The MyD88 pathway triggers IκBα degradation via IKK, enabling NF-κB subunits (e.g., NFKB1, REL) to translocate to the nucleus (75), while parallel MAPK activation phosphorylates AP-1 components (e.g., JUN, FOS) (76). These TFs collaborate with coactivators like p300 to remodel chromatin, initiating robust transcription of proinflammatory cytokines such as TNF-α, IL-6, and IL-1β (75). Simultaneously, the TRIF pathway phosphorylates IRF7, which synergizes with NF-κB and AP-1 at the IFN-β enhanceosome. This multi-protein complex recruits p300/CBP to stabilize enhancer assembly and drive IFN-β production (74, 77). Resultant IFN-β activates the expression of ISGs, including IRF1, via the JAK/STAT/IRF9 pathway. Although baseline IRF1 levels are constitutively low in resting macrophages, it integrates into the enhanceosome complex when induced, binding regulatory elements to potentiate IFN-β and ISG transcription (78). Altogether, these actions elicit an autocrine/paracrine loop where IFN-β reinforces its production, enhancing antiviral responses and solidifying M1 polarization through sustained enhancer activity. The cooperative interplay between NF-κB, AP-1, and IRFs at the enhanceosome exemplifies how innate immune signaling converges to amplify transcriptional outputs during inflammatory challenges. Remarkably, DHA, DEX, and DHA+DEX inhibited the LPS activation of NF-κB, AP-1, STAT proteins, and IRF1 (**Figure 12**).

**Figure 12.**
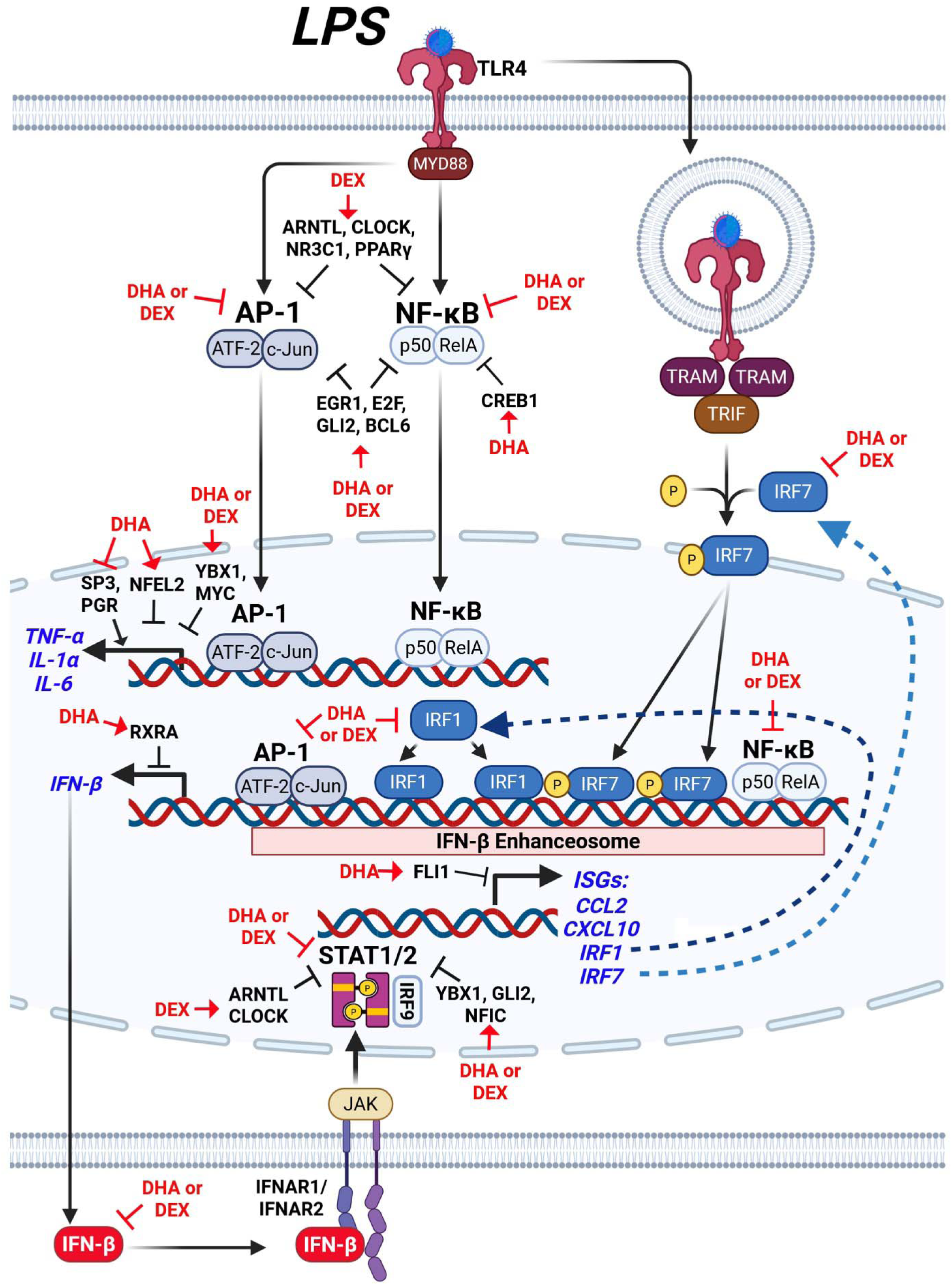
Hypothetical interplay between LPS-induced signaling pathways and the modulatory effects of DHA+DEX on type I IFN-regulated and proinflammatory gene expression. LPS stimulation activates toll-like receptor 4 (TLR4), initiating two major signaling pathways: 1) the MYD88-dependent pathway and 2) the TRIF-dependent pathway. The MYD88-dependent pathway leads to activation of the transcription factors (TFs) AP-1 (ATF-2/c-Jun) and NF-κB (p50/RelA), which induce expression of proinflammatory cytokines such as TNF-α, IL-1α, and IL-6, as well as IFN-β. Concurrently, the TRIF-dependent pathway phosphorylates IRF3/7, which forms a complex with NF-κB and AP-1, termed the IFN-β enhanceosome, to drive IFN-β production. IFN-β binds to its receptor (IFNAR1/IFNAR2), activating JAK/STAT signaling and inducing downstream ISGs such as CCL2, CXCL10, and IRF1. DHA combined with DEX suppresses the expression of inflammatory cytokines (TNF-α, IL-1α, IL-6) and IFN-β, as well as downstream genes regulated by STAT1/2, by inhibiting key IFN-β enhanceosome components. Symbols: →, increase activity; ┤, suppress activity. Created with BioRender.com.

### 4.3 DHA and DEX enrich pro-resolving regulators

The LPS-induced suppression of multiple transcriptional and post-transcriptional regulators (e.g., YBX1, EGR1, E2F, GLI2, MYC/MYCN, and NFIC) suggests that these factors form a collaborative network to suppress macrophage inflammatory signaling. Rather than acting in isolation, these regulators appear to function in a coordinated fashion, with each contributing to distinct yet overlapping mechanisms of immune suppression. For example, EGR1, in association with the NuRD complex, drives chromatin compaction and decreases accessibility at inflammatory enhancers (79), while YBX1 exerts post-transcriptional control by binding to and silencing inflammatory gene mRNAs (47, 48). This dual-layer repression system may be especially critical for maintaining stringent control over key cytokine loci, as disruption of either EGR1 or YBX1 only partially restores inflammatory responses in experimental models. Meanwhile, MYC’s role in driving glycolytic flux and stabilizing IRF4 introduces a metabolic checkpoint that intersects with NFIC’s transcriptional regulation of PTEN and SENP8, which together serve to reduce oxidative stress and further dampen immune activation (53, 55). The inhibition of GLI2 and E2F family members adds additional redundancy, with GLI2 attenuating NF-κB through Hedgehog signaling and E2F proteins modulating NF-κB dynamics and competing with AP-1 at shared promoters (49, 56). Together, these pathways could facilitate inflammatory control, where the relative importance of each regulator may depend on the cellular context or the nature of the inflammatory stimulus. Future research using gene editing approaches, such as CRISPR-mediated single and double knockouts, would clarify whether suppression of all these factors is necessary for the observed inflammatory responses or if specific pathways, such as those mediated by MYC or EGR1/YBX1, play a dominant role. The ability to dissect these interactions experimentally will not only illuminate the hierarchy of control but also inform the development of targeted anti-inflammatory therapies.

In addition to these LPS-sensitive regulators, DHA and DEX also enriched suppressive TF activities that were not markedly affected by LPS, including BCL6, NFATC2, and HNF4A. BCL6 acts as a transcriptional repressor to suppress NF-κB-driven proinflammatory genes (e.g., *IL-6*, *Ccl2*) and restrain type I IFN signaling (51, 80). At the same time, NFATC2 integrates calcium signaling, TLR4 activation, and interferon pathways to regulate macrophage immunity (81). HNF4A promotes M2 macrophage polarization via the NCOA2/GR/STAB1 axis and attenuates acute-phase gene expression, with its activation linked to improved outcomes in models of sepsis (82). The shared enrichment of these factors by DHA and DEX suggests that these therapies reinforce multiple layers of anti-inflammatory control, potentially providing a broader and more robust defense against excessive immune activation. Collectively, these findings highlight the complexity and redundancy of macrophage inflammatory regulation and point to the need for further mechanistic studies to determine which pathways are most critical and how they interact in vivo. The SLE macrophage model used here is self-renewing and genetically tractable and, therefore, amenable to CRISPR editing to address these questions.

### 4.4 DHA monotherapy selectively modulates other regulators

#### 4.4.1 Positive enrichment

DHA alone positively enriched regulatory TFs that promote anti-inflammatory and reparative functions, including NFE2L2, CEBPB, RXRA, TWIST1, STAT5A, FLI1, and FOXO1. These TFs play essential roles in resolving inflammation and maintaining macrophage homeostasis. NFE2L2 interferes with LPS-induced transcriptional upregulation of proinflammatory cytokines, including IL-6 and IL-1β, in macrophages by binding to the proximity of these genes and inhibiting RNA polymerase II recruitment (83). CEBPB is essential for M2 macrophage polarization, driving anti-inflammatory genes like *Arg1* and *Il10* via CREB-mediated induction, which is critical for resolving tissue damage (64).

RXRα (retinoid X receptor α) plays a significant role in modulating the host’s antiviral response by regulating the production of type I IFNs, particularly IFN-β (84, 85). RXRA also contributes to anti-inflammatory effects by modulating nuclear receptor-mediated gene expression networks. STAT5A activation promotes M2 macrophage polarization, favoring tissue repair and the production of anti-inflammatory cytokines (86). TWIST1 induces M2 macrophage polarization by upregulating profibrotic factors (ARG-1, CD206, IL-10, TGF-β) through direct activation of galectin-3 transcription, enhancing M2 phenotypic plasticity (62). FLI1 loss has been linked to increased IFN-regulated expression (87). FOXO1 activity is associated with both the M1 and M2 phenotypes, suggesting a more complex role in regulating macrophage polarization (67). Collectively, DHA activation of these regulatory factors might favor a reparative macrophage phenotype that counterbalances LPS-induced inflammation. Future studies could verify this mechanism by combining single-cell RNA sequencing to resolve DHA-driven transcriptional heterogeneity, time-resolved transcriptomics to map dynamic regulatory factor interactions, lipid mediator profiling to link resolvin synthesis to phenotypic outcomes, and in vivo models with genetic perturbations (e.g., RXRα/STAT5A knockdown) to test physiological relevance in LPS-challenged systems.

#### 4.4.2 Negative enrichment

DHA monotherapy also uniquely negatively enriched the activity of other proinflammatory regulators, including SP3, PGR, and RFX5. SP3 plays a critical role in promoting proinflammatory macrophage activation (M1 phenotype) by driving NF-κB-mediated transcription activation. When SP3 activity is diminished, macrophages exhibit decreased expression of M1-associated proinflammatory genes, such as *Nos2*, *Tnfa*, *Il1b*, and *Il6*, and increased expression of M2-associated anti-inflammatory markers like *Arg1* and *Retnla*. PGR, the progesterone receptor, modulates macrophage function through its activation. Stimulation of membrane-bound PGRs increases the transcription of proinflammatory genes such as *Il1b*, *Tnfa*, and *Nos2* (88), suggesting that decreased PGR activity likely suppresses these proinflammatory responses, reducing the production of inflammatory cytokines. RFX5 regulates MHC class II expression and macrophage antigen presentation (89, 90). Reduced activity of RFX5 could impair antigen presentation capacity and potentially alter cytokine signaling pathways, indirectly influencing inflammatory responses. Collectively, the suppression of these TFs by DHA likely shifts macrophage activity away from a proinflammatory phenotype.

Interestingly, DHA alone also reduced enrichment scores for suppressive factors like TCF7L2 at the 4-hr time point and CREB1, RARα, SMAD3, and GATA3 at the 8-hr time point. TCF7L2 modulates inflammatory cytokine expression in macrophages by promoting polarization toward the anti-inflammatory M2 phenotype, which suppresses proinflammatory cytokines like TNF-α and IL-6 (91). CREB1 (cAMP response element-binding protein 1) primarily suppresses proinflammatory cytokines like TNF-α and inhibits NF-κB signaling in macrophages, maintaining anti-inflammatory responses. Reduced CREB1 activity diminishes IL-10 production and decreases NF-κB suppression, amplifying proinflammatory gene transcription (92). RARα is known for its anti-inflammatory effects through modulating macrophage polarization and proinflammatory gene expression (93). SMAD3 promotes an anti-inflammatory macrophage phenotype via TGF-β signaling; its suppression increases proinflammatory cytokines such as IL-1β and TNF-α while reducing anti-inflammatory mediators like IL-10 (94). GATA3 promotes M2 differentiation and suppresses proinflammatory cytokines; reduced GATA3 activity can increase these cytokines and ISGs (95). Given their anti-inflammatory roles, decreased enrichment of these pro-resolving regulatory factors by DHA might reflect complex, time-dependent homeostatic control and suggest the need for further investigation.

### 4.5 Selective modulation of regulatory pathways by DEX monotherapy

#### 4.5.1 Positive enrichment

DEX alone selectively enriched for factors that promote resolution, including NR3C1, PPARγ, ARNTL, CLOCK, LEF1, and TFDP1. NR3C1 (glucocorticoid receptor, GR) directly inhibits proinflammatory TFs AP-1 and NF-κB, a key mechanism for suppressing inflammation (96). PPARγ reinforces these anti-inflammatory effects by inhibiting AP-1 and NF-κB, another mechanism that promotes the M2 phenotype (97). ARNTL (BMAL1) and CLOCK are circadian regulators that suppress LPS-induced proinflammatory genes (e.g., TNF-α, IL-6) in macrophages by competitively displacing NF-κB/AP-1 at enhancers and reducing H3K27ac histone acetylation, thereby limiting transcriptional activation (58, 59). ARNTL further antagonizes STAT1-mediated IFN-β signaling and stabilizes metabolic rhythms, while CLOCK reinforces these anti-inflammatory effects by curbing excessive enhancer remodeling. LEF1 expression is positively correlated with the M2 phenotype (98). Although the role of TFDP1 in inflammation remains unclear, its association with E2F factors suggests regulatory influence over immune response genes (52).

#### 4.5.2 Negative enrichment

DEX monotherapy also negatively enriched TFAP2A and ATF4, which likely dampened inflammatory pathways, as these factors regulate stress-responsive and cytokine genes (99, 100). Accordingly, DEX orchestrates a dual mechanism in macrophages by activating anti-inflammatory TFs while suppressing key proinflammatory regulators. This comprehensive modulation attenuates LPS-induced cytokine and IFN-regulated gene expression, thereby reprogramming macrophages toward an anti-inflammatory state.

### 4.6 Prior supportive evidence for O3FA and GC synergy

O3FA and GC synergy is highly consistent with previous investigations of the individual effects of these agents on TLRs, NF-κB, AP-1, STATs, and IRFs as summarized below.

#### 4.6.1 TLRs

O3FAs, notably DHA, disrupt TLR signaling through biophysical and structural mechanisms. DHA’s highly unsaturated conformation prevents stable interaction with MD2, a co-receptor for TLR4, effectively blocking TLR4 dimerization and downstream NF-κB activation (101). Beyond direct receptor interference, DHA increases membrane fluidity by incorporating phospholipid bilayers, which disperses lipid raft microdomains critical for TLR4 colocalization with CD14 (102). These biophysical effects impair receptor clustering and signaling amplification, highlighting a dual mechanism of action—direct structural inhibition and indirect membrane remodeling.

GC treatment can significantly reduce expression levels of TLR4 and MyD88 in monocytes (103). GCs also induce the expression of mitogen-activated protein kinase phosphatase-1 (MKP-1), which inhibits p38 MAPK activation downstream of TLR4, dampening cytokine production in macrophages (104). Additionally, GCs regulate TLR signaling through microRNA-mediated mechanisms, such as increasing miR-511-5p expression, which directly targets TLR4 to inhibit the production of proinflammatory cytokines (105).

#### 4.6.2 NF-κB

O3FA suppression of NF-κB is mediated through interference with both canonical and non-canonical inflammatory pathways. DHA and EPA competitively inhibit arachidonic acid metabolism, reducing proinflammatory prostaglandin E2 and leukotriene B4 production, which are known to enhance NF-κB activity (106). Oxidized metabolites of O3FAs, such as 18-HEPE and 17-HDHA, exhibit enhanced potency by activating PPARα, which sequesters NF-κB coactivators and promotes its nuclear export (107, 108). O3FAs also directly inhibit IκB kinase (IKK) phosphorylation, preventing IκB degradation and NF-κB nuclear translocation in macrophages (109).

GC interference with NF-κB signaling is multifaceted and central to its anti-inflammatory effects. GCs induce the synthesis of IκBα, which sequesters NF-κB in inactive cytoplasmic complexes, preventing its nuclear translocation and transcriptional activity (110). The GC receptor (GR) physically associates with the p65 subunit of NF-κB, disrupting its DNA-binding and transcriptional activation capabilities (54). GCs also induce the expression of GC-induced leucine zipper (GILZ), which binds to the transactivation domain of activated NF-κB p65, further inhibiting its activity (111).

#### 4.6.3 AP-1

O3FAs attenuate AP-1 signaling by targeting MAPK cascades. In murine macrophages, O3FAs suppress p44/42 and JNK/SAPK phosphorylation—steps preceding AP-1 activation—lead to reduced AP-1 activity and subsequent downregulation of proinflammatory cytokine genes in macrophages, confirming transcriptional-level anti-inflammatory effects (112). EPA suppresses phosphorylation of p38 MAPK and JNK/SAPK in human monocytic THP-1 cells, reducing c-Fos/c-Jun heterodimer formation and AP-1 DNA-binding activity (113). O3FA inhibition of AP-1 may be linked to the upregulation of MAPK phosphatase-1 (MKP-1), which dephosphorylates and inactivates JNK (114).

GC-mediated suppression of AP-1 signaling occurs through several mechanisms. GRs physically interact with c-Jun and c-Fos, components of AP-1, inhibiting their transcriptional activity without requiring GR binding to DNA (115, 116). GCs also inhibit the phosphorylation and activation of JNK, thereby reducing AP-1 activity (117, 118). The induction of MAPK phosphatase-1 (MKP-1) by GR activation further suppresses AP-1 activity by dephosphorylating and inactivating JNK (119).

#### 4.6.4 STATs

O3FAs and GCs also attenuate STAT activation. We have previously found that DHA inhibits the expression of STAT1/STAT2-target genes in LPS-treated macrophages (31). RvD2, a pro-resolving metabolite of DHA, suppresses the phosphorylation of STAT1 in bone marrow-derived macrophages (120). DHA and its metabolites also inhibit STAT3 phosphorylation in cancer cells (121–124).

GCs primarily inhibit STAT1 through the induction of SOCS1, which inhibits STAT1 activation by degrading phosphorylated JAK2 (73). Furthermore, GCs suppress TLR-mediated STAT1 phosphorylation at Ser727 and Tyr701 during later phases of activation, impairing its nuclear translocation and transcriptional activity. Recruitment of GR to DNA-bound STAT3 is associated with trans-repression or transcriptional antagonism (125).

#### 4.6.4 IFN signaling

O3FAs indirectly regulate IRFs and IFNAR signaling by inhibiting ISG expression in LPS-treated macrophages (31). DHA also attenuates IFNAR signaling by reducing STAT1 phosphorylation, thereby blunting IFN-driven inflammatory gene expression (126). Meanwhile, GCs have been shown to interfere with IRF signaling by suppressing STAT1 mRNA transcription, leading to reduced activation of IRF-dependent pathways and diminished IFN-inducible gene expression, particularly in macrophages (127). DEX inhibits IRF3 phosphorylation and nuclear translocation in macrophages by suppressing TBK1, a kinase crucial for IRF3 activation (128). The GC receptor sequesters GRIP1, a coactivator for IRF3 and IRF9, preventing their transcriptional activity and disrupting the activation of ISGs (71). GCs antagonize the co-recruitment of IRF3 and NF-κB subunit p65 to ISRE-containing promoters, thereby reducing ISG transcription (71). GCs interfere with IFN receptor signaling by inhibiting the assembly of the STAT1-STAT2-IRF9 (ISGF3) transcription complex, essential for type I IFN signaling, and also prevent the nuclear translocation of IRF9, a critical step for triggering IFN-responsive gene expression (129). Furthermore, GCs block IFN-induced IRF1 mRNA levels, disrupting transcriptional activation of interferon-responsive genes regulated by IRF elements in the GC receptor promoter region (130). These findings suggest a strong synergistic potential between O3FAs and GCs that should continue to be investigated in future *in vitro* and *in vivo* studies.

### 4.7 Limitations

Although we present herein compelling evidence for O3FA+GC synergy in inhibiting LPS-induced inflammation in SLE macrophages, we acknowledge that our study has limitations. First, our use of an *in vitro* LPS-activated NZBWF1 macrophage model, while mechanistically informative, does not capture the full complexity of human SLE, as it lacks multicellular interactions and the tissue-specific microenvironment present in an *in vivo* model. Second, our analysis was limited to early transcriptional responses (4-8 hr post-LPS treatment), creating uncertainty about whether the observed DHA and DEX synergy is sustained or subject to rebound effects during chronic inflammation. Third, although key transcriptional regulators were identified by functional enrichment, the precise molecular mechanisms underlying the observed synergy, such as direct receptor crosstalk, epigenetic changes, or metabolic reprogramming, remain uncharacterized. Fourth, the efficacy and safety of low-dose DEX and DHA cotreatment have not yet been validated in preclinical SLE models or clinical settings, where factors such as bioavailability, off-target effects, and patient heterogeneity could significantly impact therapeutic outcomes. These limitations highlight the need for further mechanistic, longitudinal, and preclinical studies to advance this promising combinatorial approach toward clinical application.

## 5 Conclusions

The findings presented herein demonstrate that low-dose DHA synergizes with low-dose DEX to suppress LPS-induced inflammatory signaling in SLE-modeled macrophages potently, amplifying DEX’s efficacy by 10- to 100-fold in attenuating ISGs. Computational synergy analysis revealed robust cooperativity between DHA and DEX at nanomolar-to-micromolar concentrations, with RNA-seq profiling showing that the combination induces distinct transcriptional programs, suppressing proinflammatory pathways (NF-κB, STATs) while upregulating resolution-associated regulators (TWIST1, BCL6) and metabolic shifts favoring M2 polarization. This dual targeting of shared (AP-1, IRF1) and unique (YBX1, EGR1) transcriptional nodes enables the combinatorial regimen to override LPS-driven hyperinflammation more effectively than either agent alone. The ability of DHA to enhance DEX’s suppressive potency at subtherapeutic doses suggests a feasible strategy to mitigate GC toxicity while maintaining efficacy in SLE management. These data position O3FA-GC co-therapy as a mechanistically grounded approach to recalibrate macrophage phenotypes and reduce steroid reliance in the management of autoimmune hyperinflammation.

## 7 Author Contributions

LH: study design, data analysis/interpretation, figure preparation, manuscript preparation; RN: data curation, figure preparation, manuscript preparation; JJ: data analysis; AA: study design; JH: study design, project funding, manuscript preparation; AO: study design, project funding, manuscript editing; JP: study design, project funding, manuscript preparation; OM: FLAM isolation, data analysis/interpretation, figure preparation, manuscript preparation.

## 9 Acknowledgments

The authors thank the Michigan State University Genomics Core Facility for their expertise and assistance with sequencing all RNA-seq samples.

## 10 Data Availability Statement

RNA-seq datasets are deposited in the Gene Expression Omnibus (GEO) with the accession identifier GSE298247.

