## Supplemental Material for "Omega-3 Fatty Acid Synergy with Glucocorticoid in Lupus Macrophages: Targeting Pathogenic Pathways to Reduce Steroid Dependence"

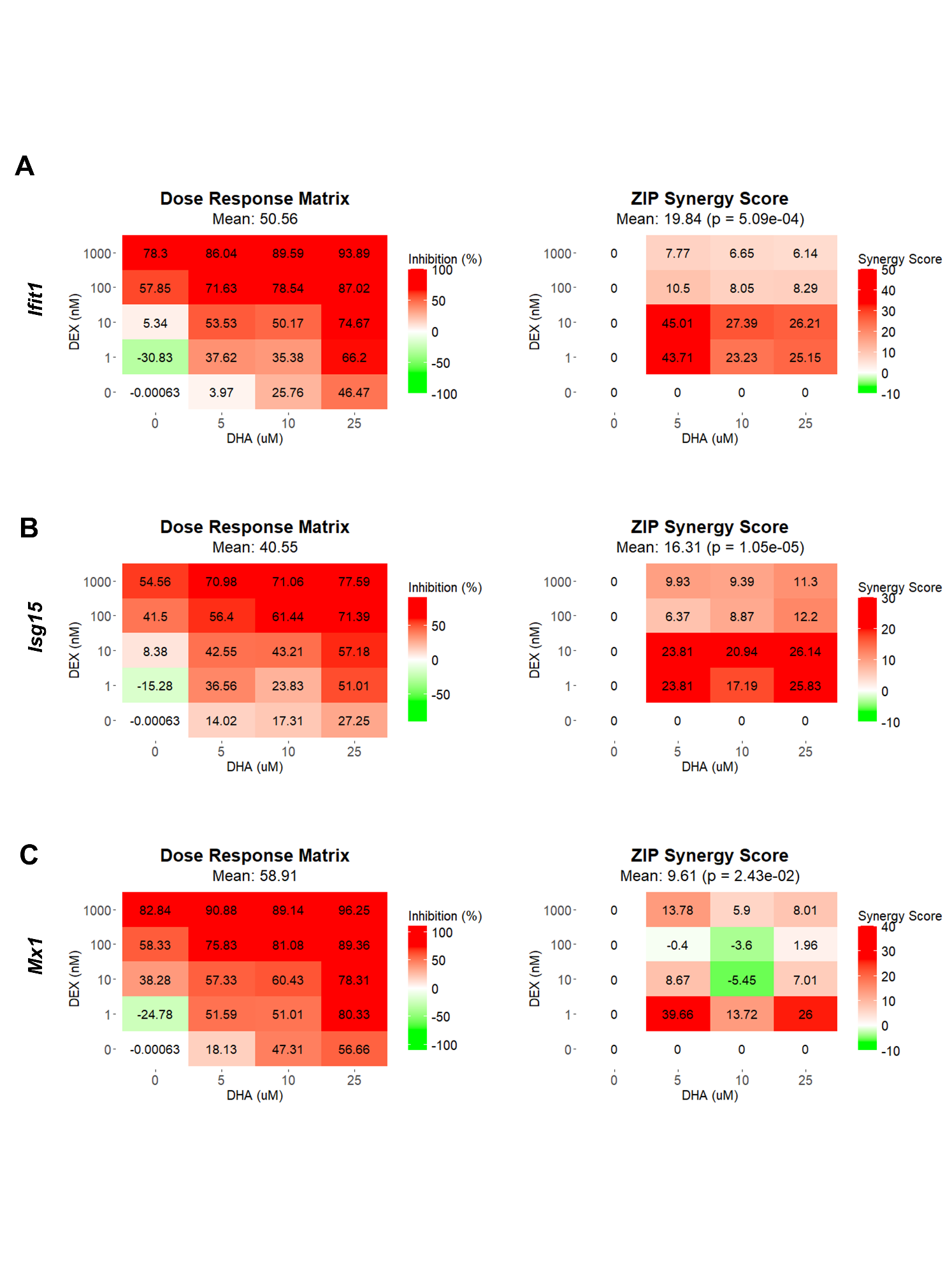

**Supplementary Figure 1. DHA and DEX synergistically inhibit the expression of IFN-stimulated genes.** *Ifit1* **(A)**, *Isg15* **(B)**, and *Mx1* **(C)** were measured by qRT-PCR in FLAMs stimulated with LPS (20 ng/mL) for 4 hr. Cells were pretreated with either VEH containing no DHA or RPMI media containing 25 µM, 10 µM, or 5 µM DHA at -24 hr. Cells were then treated with VEH containing no DEX or varying concentrations of DEX (1 nM-1 µM) -1 hr prior to LPS treatment. SynergyFinder version 3.14.0 was used to generate inhibition matrices and ZIP synergy matrices for each gene. Inhibition matrices show the average of 3 experimental replicates. Individual and mean ZIP synergy scores were calculated using an average of 3 experimental replicates. Synergy score > 0, synergistic interaction; synergy score = 0, additive effect; synergy score < 0, antagonistic interaction.

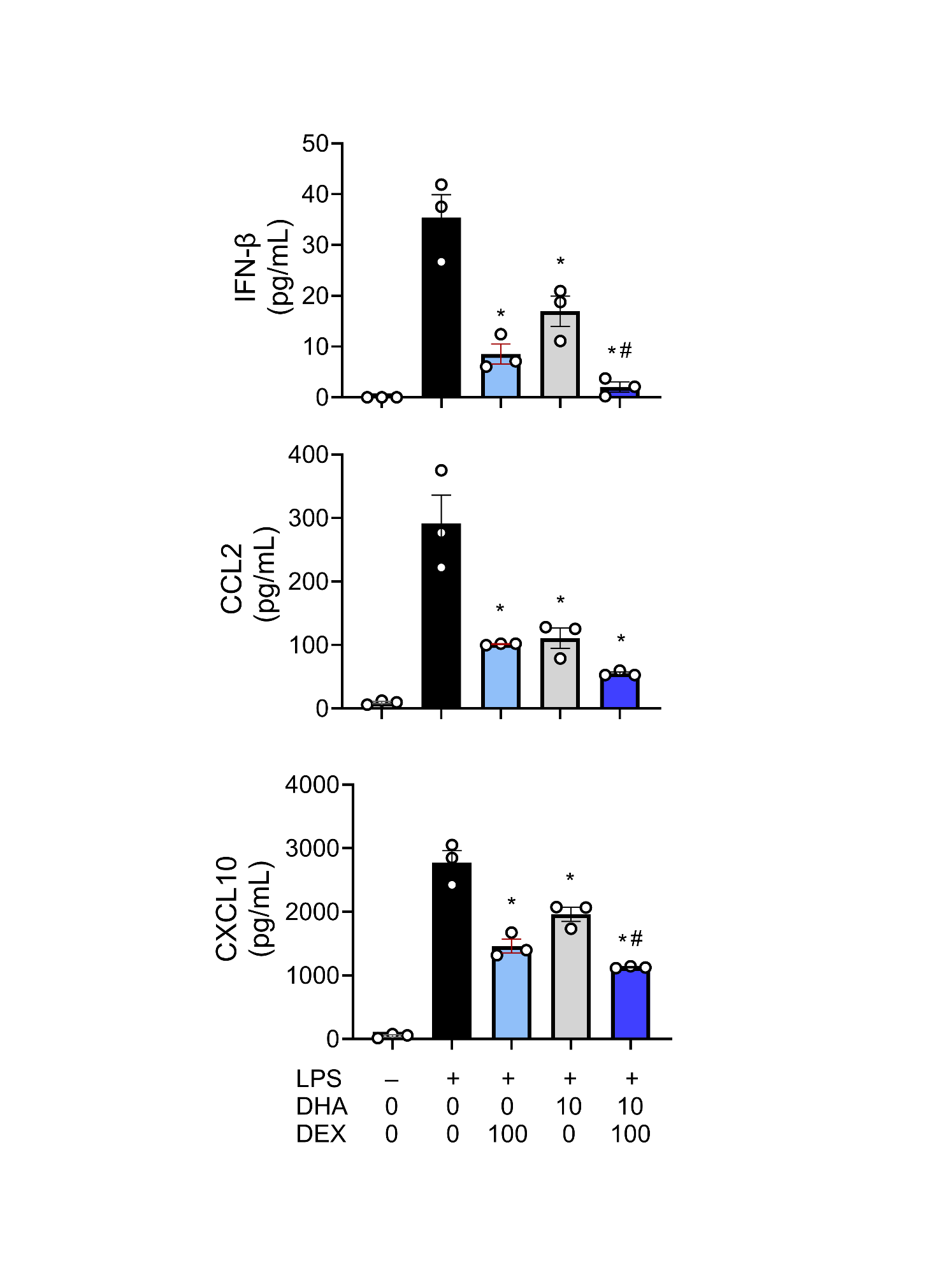

**Supplementary Figure 2. DHA+DEX combination treatment suppresses type I IFN-related protein secretion.** Cells were pretreated with either VEH containing no DHA or RPMI media containing 10 µM or 25 µM DHA at -24 hr. Cells were then treated with VEH containing no DEX or RPMI media containing 100 nM or 1000 nM DEX -1 hr prior to LPS treatment. Type I IFN-related proteins (i.e., IFN-β, CCL2, CXCL10) were measured by ELISA in supernatants from FLAMs stimulated with LPS (20 ng/mL) for 24 hr. Data are shown as mean ± SEM. n=3 biological replicates. p<0.05; *Significant compared to LPS/VEH; #Significant compared to DHA alone; †Significant compared to DEX alone.
